# RNA transcripts suppress G-quadruplex structures through G-loop formation

**DOI:** 10.1101/2023.03.09.531892

**Authors:** Koichi Sato, Aiko G.M. Hendrikx, Puck Knipscheer

**Author notes:** Correspondence can be addressed to and.

## Abstract

The eukaryotic genome contains numerous guanine-rich sequences that regulate DNA replication and transcription by folding into non-canonical structures called G-quadruplexes (G4s)^1^. Compromised G4 resolution causes genomic instability and diseases linked to cancer susceptibility and aging^2^. However, how G4s are resolved is poorly understood. G4 structures are found in genomic regions containing DNA:RNA hybrids, also referred to as R-loops^3, 4^. When an R-loop and G4 structure form exactly opposite to each other, a G-loop structure is formed. G-loops have been observed directly in prokaryotes^5^ but their function is unknown. Using the *Xenopus* egg extract system, we show that G-loop structures act in suppressing mutagenic G4 structures thereby preventing genomic instability. Mechanistically, the RAD51 recombinase binds to the G4-opposing strand and promotes hybridization of RNA transcripts supplied by the hnRNPA1 ribonucleoprotein complex independently of ongoing transcription. The resulting G-loop then induces FANCI-FANCD2 monoubiquitination that triggers site-specific incision of the hybrid strand by the SLX4-XPF-ERCC1 complex. G-loop incision requires prior G4 unwinding by the DHX36 and FANCJ helicases and allows immediate DNA synthesis past the G4 motif. This resynthesis step impacts local chromatin states. This work establishes a G-loop-dependent mechanism that prevents mutagenic consequences of both G4 and R-loop structures.

---

G4 structures are prevalent in promotors, enhancers, and telomeres where they play a role in transcriptional regulation and telomere maintenance^6, 7^. Nevertheless, they must also be timely suppressed by unwinding throughout the cell cycle to maintain genome integrity^8, 9^. G4 structure unfolding requires specialized DNA helicases, defects in which cause various human genetic and age-related diseases characterized by telomere instability and predisposition to cancers^2^. Because G4 structures block DNA replication^10^, G4 resolution in S phase is crucial. We previously showed that during DNA replication G4 structures are resolved through sequential action of two G4 helicases, DHX36 and FANCJ and subsequent DNA replication past the G4 motifs^10^ (Figure S1a). However, how G4 structures are suppressed outside DNA replication to regulate their biological roles and avoid DNA damage^11, 12^ remains a key question. While current models envision that G4 resolution is accomplished by simply unwinding and reannealing of the two DNA strands, it is not clear whether additional regulatory steps and proteins are involved. We therefore set out to study DNA replication-independent G4 resolution mechanisms using the cell-free *Xenopus* egg extract system. For this, we used a pG4^BOT^ plasmid that contains a canonical G4 structure (G4^G3N^) on one strand and a non-complementary sequence opposite the G4 motif (pG4^BOT^, Figure S1b). This bubble structure was necessary to allow stable G4 structure formation^10^ and resembles a physiological situation where G4 formation will result in displacement of the opposite strand. We incubated pG4^BOT^ in high-speed supernatant (HSS) and subsequently added nucleoplasmic extract (NPE) in the presence or absence of Geminin that blocks replication initiation. Products were digested with NotI, that excises an 81-bp fragment containing the G4 region, and ClaI, that specifically incises the non-G4 strand only when DNA synthesis occurs on the strand (Figure S1c). Interestingly, in the absence of DNA replication, incubation of pG4^BOT^ yielded, in addition to the bubble region, two faster-migrating products upon separation by native PAGE (Figure S1d). These products corresponded to the duplex form of the G4- and non-G4 strands (81-bp and 51-bp fragments, respectively), as replication of pG4^BOT^ resulted in the same molecules (Figure S1d). Notably, in the presence of Geminin, the 81-bp product preferentially accumulated and this was more prominently observed in a mitotic *Xenopus* egg extract, and in diluted NPE only, both extracts that don’t support DNA replication (Figure S1e,f). In contrast, in HSS only, or in the absence of a G4 structure, the enhanced accumulation of the 81-bp fragment was lost (Figure 1a, Figure S1f). This suggests that a G4 suppression mechanism that converts the G4-containing strand to dsDNA is active under specific conditions (Figure S1c). This mechanism is not G4^G3N^ specific but also acts on other G4 structures since similar results were obtained with pTEL^BOT^, containing a telomeric G4 (Figure S1b,g). Stabilization of the structure by the G4 ligand PhenDC_3_ prevented conversion of the G4-containing strand to dsDNA, indicating that conversion requires G4 structure unfolding (Figure 1a). Moreover, we found the same G4 helicases to be involved in unwinding compared to the replication-dependent mechanism, because replication-independent G4 conversion was also fully blocked by double depletion of FANCJ and DHX36. This defect was restored by addition of wild-type but not the ATPase-dead mutant proteins (Figure S1h-j).

**Figure 1.**
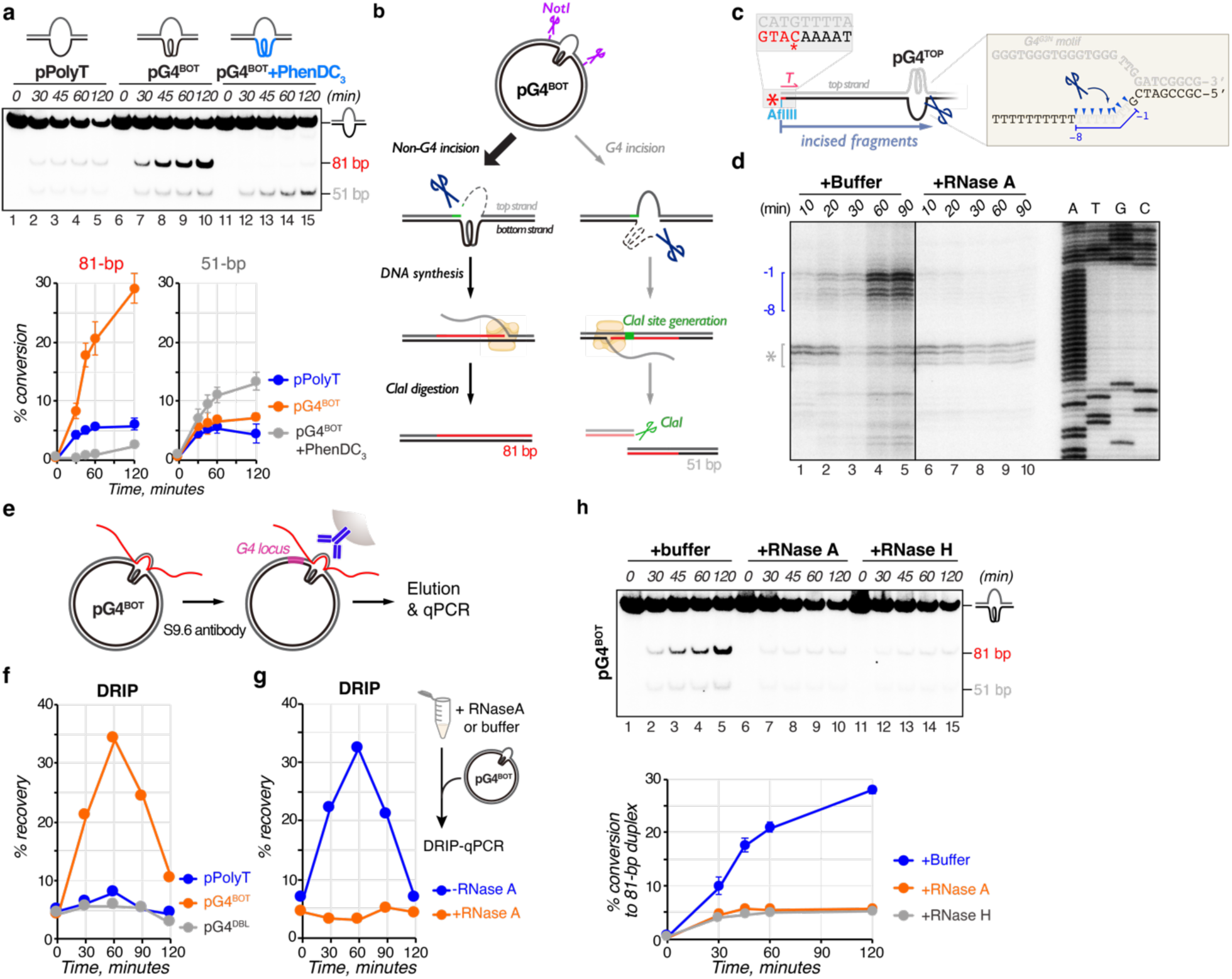
G-quadruplex suppression through RNA invasion and nucleolytic incisions. **a**, pPolyT and pG4^BOT^ were incubated in NPE in the presence or absence of PhenDC_3_. At various times, DNA was isolated, digested with NotI and ClaI, end-labeled with ^32^P-α-dCTP, separated by native PAGE, and visualized by autoradiography (top). The 81-bp and 51-bp fragments were quantified and the conversion percentage was calculated by comparing their intensity to the total intensity of all fragments at time point 0. Conversion (%) was plotted against time with standard deviations (bottom, n=3). **b**, Model to explain how the G4-forming region is converted to complementary dsDNA in pG4^BOT^. **c,** Schematic representation of incision products generated when pG4^TOP^ is incubated in NPE, products are digested with AflIII, and end-labeled with ^32^P-α-dCTP (asterisk)(left). Region surrounding the incision sites is enlarged and incision positions observed in **d** are indicated (blue arrow heads) (right). Incision products are numbered −8 to −1, where the −1 product corresponds to a product in which incision takes place between the first and the second nucleotide upstream of the ss/dsDNA junction. **d**, pG4^TOP^ was incubated in NPE that was pre-treated with RNaseA or buffer. At various times DNA was isolated, digested with AflIII, end-labelled with ^32^P-α-dCTP, separated by denaturing PAGE, and visualized by autoradiography. A sequencing ladder derived from extension of primer *T* annealed to pG4^TOP^ (*T* in **c**) was used as size markers. Incision products are indicated with a blue bracket. The asterisk indicates products generated by star activity of AflIII. **e,** Schematic of the DNA-RNA immunoprecipitation (DRIP) assay. G4-containing plasmids are incubated in NPE, products are isolated and immunoprecipitated with the S9.6 antibody (blue). The co-precipitated DNA is amplified by quantitative PCR (qPCR) with primers specific to the G4 locus (magenta). **f,** pPolyT, pG4^BOT^, and pG4^DBL^ were incubated in NPE, and products were analysed by DRIP-qPCR with primers for the G4 locus. Relative values compared to input signals were plotted. **g,** pG4^BOT^ was incubated in NPE that was pre-treated with RNaseA or buffer and products were analysed as in **f**. **h**, pG4^BOT^ was incubated in NPE that was pre-treated with RNaseA, RNaseH, or buffer, and products were analysed as in **a** (n=3).

## G4-dependent nucleolytic incision of the non-G4 strand

G4 suppression requires DNA synthesis past the G4 motif (Figure 1b). In agreement with this, labeled nucleotides were preferentially incorporated into a 270 bp-fragment containing the G4 in both pG4^BOT^ and pTEL^BOT^ upon incubation in NPE (Figure S2a-c). This could be reminiscent of the DNA repair synthesis that specifically colocalizes with G4 sequences in post-mitotic neurons^13^. Because DNA synthesis requires a 3’ DNA end for initiation, the plasmids must be locally incised (Figure 1b). To detect nucleolytic incisions upon incubation in NPE, we linearized the plasmid with AflIII, end-labeled the products, and separated them on a sequencing gel (Figure 1c). When G4^G3N^ was placed on the top strand (pG4^TOP^), a cluster of fragments accumulated that corresponds to incisions 1 to 8 nucleotides upstream of the ss/dsDNA junction (−1 to −8 position) (Figure 1c,d). Similar incision products were observed for pTEL^TOP^ (Figure S2d) but only when the G4 was placed on the top strand (Figure S2e). Therefore, G4 structures induce nucleolytic incisions selectively on the non-G4 strand, allowing DNA synthesis past the G4 (Figure 1b).

## G-loop formation triggers G4 processing

The major increase in nucleolytic incisions only occurred between 30 and 60 minutes, suggesting that they are preceded by other essential step(s). Previous studies suggest that G4 sequences overlap with R-loop hotspots genome wide^3, 4^, although the formation of DNA:RNA hybrid across from a G4 structure, called ‘G-loop’^5^, has only been directly observed in bacteria. We thus asked whether RNA invades into the G4 region to promote nucleolytic incision. After incubation in NPE, we isolated the plasmids and assessed R-loops using DNA:RNA immunoprecipitation followed by qPCR (DRIP-qPCR) (Figure 1e). Strikingly, pG4^BOT^ exhibited a marked R-loop signal that peaked at 60 minutes and declined to the basal level by 120 minutes, comparable to the half-life time of R-loop in human cells^14^, while the signal was hardly detected in the absence of the G4 structure (Figure 1f). R-loops were specifically detected at the G4 region and formed with similar kinetics in pTEL^BOT^ (Figure S3a,b). R-loop signal was lost upon pre-treatment with RNaseH, an RNase that specifically degrades DNA:RNA hybrids, and on plasmids carrying G4s on both strands (pG4^DBL^) or in the context of dsDNA (pdsG4^BOT^), suggesting G4-dependent DNA:RNA hybrid formation on the non-G4 strand (Figure 1f, Figure S3c-e). These results are consistent with observations in human cells that G4 stabilization triggers immediate and global increase in R-loops^3^. The invading RNA is most likely derived from oocyte transcripts (Figure S3f), because pre-treatment with RNaseA drastically reduced R-loop levels, while depletion of RNA polymerase II and treatment with `α`-amanitin, an inhibitor of eukaryotic RNA polymerases, had no effect (Figure 1g, Figure S3g-i). Collectively, RNA transcripts invade into the non-G4 strand across from a G4 structure independently of ongoing transcription, forming a G-loop.

To determine the functional relevance of G-loop formation, we assessed nucleolytic incisions in the presence of RNaseA. This treatment severely reduced the cluster of incisions for both G4^TEL^ and G4^G3N^ (Figure 1d, Figure S2d). Moreover, DNA synthesis past the G4 and G4 conversion was disrupted in the presence of RNaseA or RNaseH (Figure 1h, Figure S2b,c, Figure S3j). Notably, the bubble structure underwent marked degradation in the absence of, or upon inhibition of G-loop formation (Figure 1a,h, Figure S1g, Figure S3j), indicating a protective role of DNA:RNA hybrids from nuclease attacks. Together, formation of G-loops is prerequisite for G4 suppression.

## RAD51 promotes G-loop formation

R-loop formation in *trans*, in which the RNA is not supplied co-transcriptionally, can be mediated by the RAD51 recombinase in yeast and mammalian cells^15, 16^. To examine a role for RAD51 in G-loop formation, we depleted RAD51 and BRCA2 and found that both depletions prevented R-loop accumulation in pG4^BOT^ (Figure 2a,b). G4 suppression was also severely prevented by RAD51 depletion (Figure S4a-c). While co-depletion of BRCA2 with RAD51 complicated rescue experiments (Figure 2a, Figure S4a), the defect was partially restored by pre-loading wild-type RAD51 onto the plasmid but not by pre-loading the homologous pairing-defective RAD51^T128R^ mutant^17^ (Figure S4a-d). These data suggest that RAD51 loading onto the plasmid by BRCA2 promotes G-loop formation. In agreement with this, plasmid-based chromatin immunoprecipitation (ChIP) detected RAD51 and BRCA2 specifically at the G4 region of pG4^BOT^ immediately after reaction initiation, and both proteins mostly disappeared from this region by 60 minutes when G-loop formation peaked (Figure 2c,d). Notably, RNaseA treatment did not affect the levels of RAD51 and BRCA2 at the G4 region, indicating that they accumulate prior to homology search (Figure S4e). Recruitment of BRCA2 and RAD51 was lost when a G4 was present on both strands (Figure 2d). Thus, these proteins are loaded onto the non-G4 strand to promote DNA:RNA hybridization through homologous pairing.

**Figure 2.**
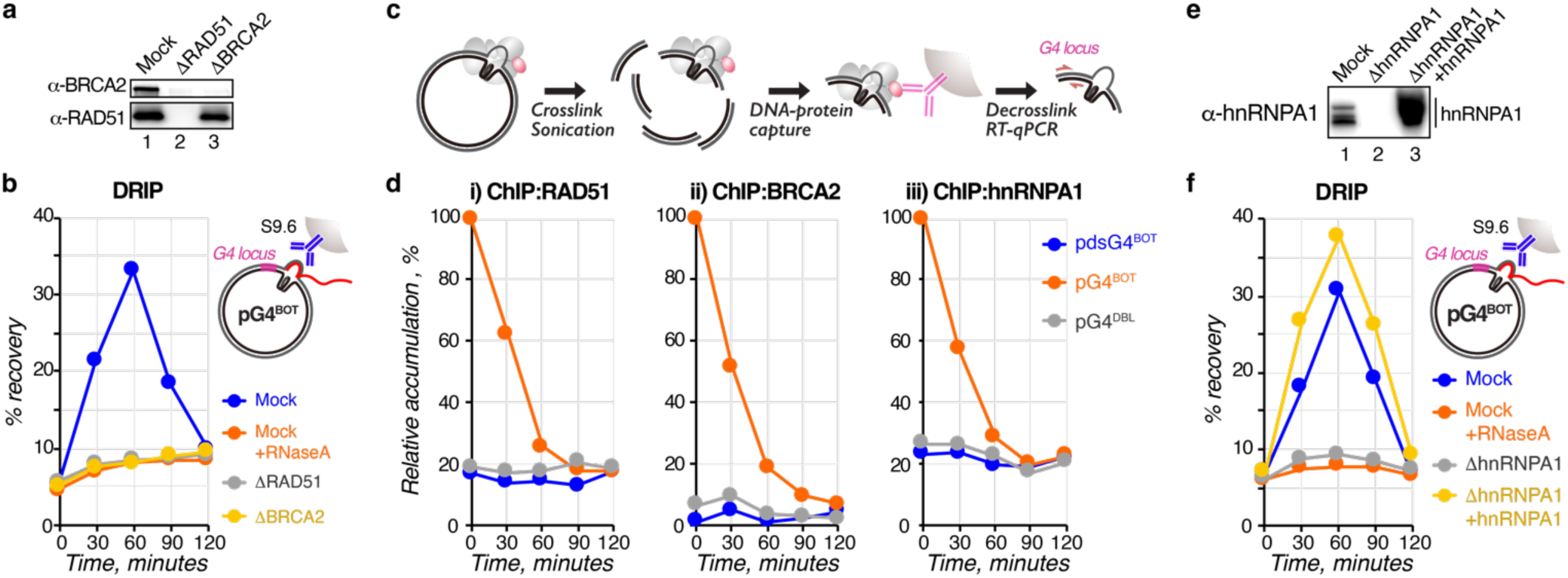
Mechanism of G-loop formation by RAD51, BRCA2 and the hnRNPA1 complex. **a,** Mock-, RAD51-, and BRCA2-depleted NPEs were analysed by Western blot with RAD51 and BRCA2 antibodies. **b,** pG4^BOT^ was incubated in the NPEs described in **a**, and products were analysed by DRIP-qPCR using primers for the G4 locus. Where indicated, NPE was pre-treated with RNaseA. Relative values compared to input signals were plotted. **c**, Scheme of the chromatin immunoprecipitation (ChIP) assay. G4-containing plasmids are incubated in NPE, DNA and proteins (ovals) are crosslinked, sonicated, and immunoprecipitated with the antibody of interest (magenta). The co-precipitated DNA is amplified by quantitative PCR (qPCR) with primers (magenta arrows) specific to the G4 locus. **d,** pdsG4^BOT^, pG4^BOT^, and pG4^DBL^ were incubated in NPE, and products were analysed by ChIP-qPCR with RAD51, BRCA2, and hnRNPA1 antibodies and a primer pair for the G4 locus. The relative values compared to the highest signal among the conditions were plotted. **e,** Mock- and hnRNPA1-depleted NPEs supplemented with buffer or the hnRNPA1 complex purified from extract, were analysed by Western blot with the hnRNPA1 antibody. **f,** pG4^BOT^ was incubated in the NPEs described in **e**, and products were analysed by DRIP-qPCR as in **b**. Where indicated, NPE was pre-treated with RNaseA.

## The hnRNPA1 complex shuttles RNA transcripts to G4s

We then asked how the RAD51 nucleoprotein filament engages with endogenous RNAs to promote G-loop formation. RNA transcripts are bound by heterogeneous nuclear ribonucleoproteins (hnRNPs)^18^. One of these, hnRNPA1, colocalizes with R-loops and repair synthesis hotspots^13, 19^, interacts with RNA transcripts that are involved in R-loop formation *in trans*^15, 20^, and causes genome instability at G4-forming loci when mutated^21, 22^. Depletion of hnRNPA1 from extract strongly reduced R-loop formation on pG4^BOT^, and this defect was reversed by addition of the hnRNPA1 complex purified from NPE (Figure 2e,f, Figure S4d). Furthermore, hnRNPA1 depletion abrogated G4 conversion which was also largely rescued by hnRNPA1 purified from NPE (Figure S4f,g). Recruitment of hnRNPA1 to the G4 region of pG4^BOT^ occurred with similar kinetics to RAD51 and BRCA2 but no recruitment was observed on pG4^DBL^ or pdsG4^BOT^ (Figure 2d), indicating that hnRNPA1 is also loaded onto the non-G4 strand. RAD51 and hnRNPA1 recruitment was independent of each other but depletion of RAD51 resulted in persistence of hnRNPA1 at the G4 region and vice versa (Figure S4h,i). Together, these data support a model in which the hnRNPA1 complex recruits RNA transcripts to G4s to promote G-loop formation mediated by RAD51.

## The FANCI-FANCD2 complex recruits SLX4-XPF-ERCC1 to promote incisions

R-loops colocalize with a key DNA repair complex of the Fanconi anemia pathway, FANCI-FANCD2 (ID) in cells^23, 24^, but the functional relevance of this is largely unknown. To examine this, we monitored FANCD2 recruitment and found that it specifically accumulated at the G4^G3N^ region with similar kinetics to G-loop formation (Figure 3a). Consistent with FANCD2 monoubiquitination being required for R-loop retention^24^, robust FANCD2 monoubiquitination was detected upon incubation of both pG4^BOT^ and pTEL^BOT^ in extract (Figure S5a). Inhibition of G-loop formation by RNaseA treatment or hnRNPA1 depletion attenuated FANCD2 ubiquitination and blocked FANCD2 loading to the G4 region (Figure 3a, Figure S5a,b). Moreover, when FANCD2 ubiquitination was prevented by depletion of the FA core E3 ligase subunit FANCA, FANCD2 was no longer recruited even though G-loops still formed (Figure S5c-g). These data show that G-loops are required to recruit the ID complex by stimulating FANCD2 monoubiquitination.

**Figure 3.**
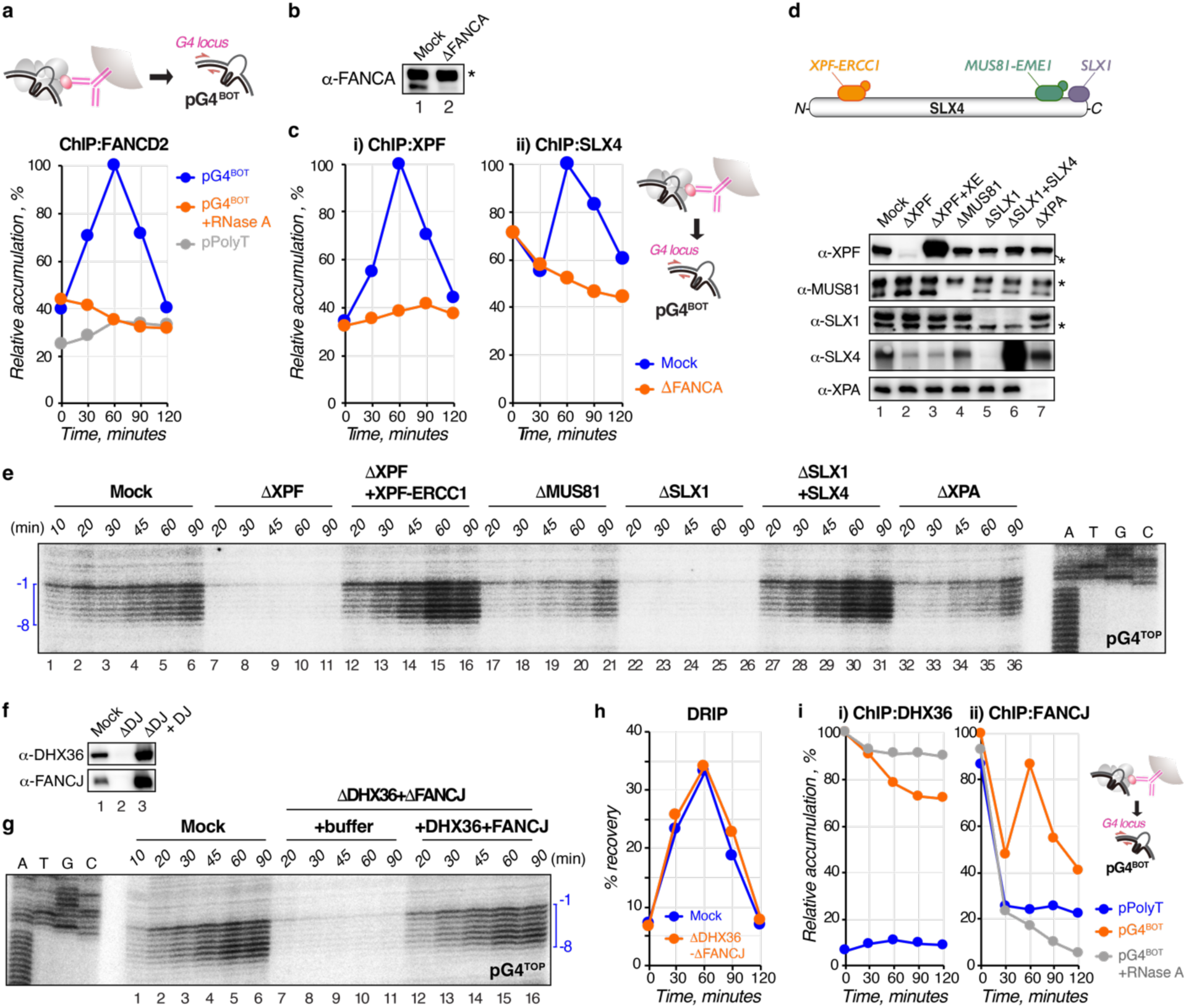
Mechanism of nucleolytic incision during G-quadruplex suppression. **a**, pG4 ^BOT^ and pPolyT were incubated in NPE pre-treated with buffer or RNaseA, and products were analysed by ChIP-qPCR with the FANCD2 antibody and a primer pair for the G4 locus. The relative values compared to the highest signal among the conditions were plotted. **b**, Mock- and FANCA-depleted NPEs were analysed by Western blot with the FANCA antibody. The asterisk represents a non-specific band. **c**, pG4 ^BOT^ was incubated in the NPEs as described in **b**, and products were analysed by ChIP-qPCR as in **a** with XPF (i) and SLX4 (ii) antibodies using primers for the G4 locus (schematic, right). **d**, Mock-, XPF-, MUS81-, SLX1-, and XPA-depleted NPEs supplemented with buffer, or where indicated with the XPF-ERCC1 complex or SLX4, were analysed by Western blot with XPF, MUS81, SLX1, SLX4, and XPA antibodies (bottom). The asterisks represent non-specific bands. Schematic representation of SLX4 and its interacting nucleases is depicted (top). **e**, pG4^TOP^ was incubated in the NPEs as described in **d**, and products were digested with AflIII, end-labelled, separated by denaturing PAGE alongside a sequencing ladder, and visualized by autoradiography. Incised fragments (−1 to −8) are indicated with a bracket. **f**, Mock- and DHX36-FANCJ-depleted NPEs supplemented with buffer or wild-type DHX36 and FANCJ were analysed by Western blot with DHX36 and FANCJ antibodies. **g,** pG4^TOP^ was incubated in the NPEs as described in **f,** and products were analysed by denaturing PAGE as in **e**. **h,** pG4^BOT^ was incubated in the NPEs as described in **f**, and products were analysed by DRIP- qPCR using primers for the G4 locus. Relative values compared to input signals were plotted. **i,** pPolyT and pG4 ^BOT^ were incubated in NPE, and products were analysed by ChIP-qPCR with DHX36 (i) and FANCJ (ii) antibodies using primers for the G4 locus (schematic, right). Where indicated, NPE was pre-treated with RNase A. The relative values compared to the highest signal among the conditions were plotted.

Defective FANCD2 ubiquitination blocks DNA interstrand crosslink (ICL) repair and is associated with Fanconi anemia, a cancer predisposition syndrome characterized by bone marrow failure^25^. During ICL repair, the ubiquitinated ID complex promotes recruitment of the SLX4-XPF-ERCC1 endonuclease complex required for ICL unhooking^26^. We therefore wondered whether an analogous mechanism acts in G-loop processing. Using ChIP, we found that both XPF and SLX4 were recruited to the G4^G3N^ region, peaking at 60 minutes similar to FANCD2, and this was dependent on FANCD2 ubiquitination (Figure 3b,c). In addition, SLX4 depletion prevented XPF, but not FANCD2 recruitment (Figure S5h,i), indicating that XPF and SLX4 are recruited as a complex after the ubiquitinated ID complex. This suggests that the SLX4-XPF-ERCC1 complex might be involved in nucleolytic incisions on the non-G4 strand. We confirmed this by depletion of XPF-ERCC1 which abrogated the incisions in both pG4^TOP^ and pTEL^TOP^, which was fully rescued by addition of recombinant XPF-ERCC1 (Figure 3d,e, lanes 7-16, Figure S5j,k). The other SLX4 interacting nucleases, SLX1 and MUS81, as well as XPA, the essential localizer of XPF-ERCC1 during nucleotide excision repair, were not required for these incisions (Figure 3d,e, Figure S5k). Together, the monoubiquitinated ID complex promotes site-specific incisions on the non-G4 strand by recruiting the SLX4-XPF-ERCC1 nuclease complex.

## Nucleolytic incision requires G4 unwinding

The XPF-ERCC1 complex preferentially acts on ds/ssDNA junctions with splayed ssDNA arms^27^. However, in G-loops, the 5’ ssDNA forms a G4 structure, suggesting that resolution of the structure may be required for the incision. To prevent G4 unwinding, we double-depleted FANCJ and DHX36 and found that it abrogated the incision on the non-G4 strand of pG4^TOP^, which was fully restored by addition of both recombinant proteins (Figure 3f,g). In contrast, the double depletion had no effect on R-loop formation (Figure 3h), indicating that the incision defect is not due to compromised G-loop formation. Therefore, G4 resolution is pre-requisite for incisions on the non-G4 strand.

Similar to the mechanism we previously described for DNA replication-dependent G4 resolution^10^ (Figure S1a), DHX36 and FANCJ seem to act in a consecutive fashion during G-loop formation. DHX36 specifically accumulated at the G4^G3N^ region at the start of the reaction, after which the DHX36 signals gradually dropped by ∼30%, similar to the degree of G4 conversion we observed (Figure 3i). In contrast, FANCJ accumulated at the G4 region 60 minutes after the start of the reaction concomitant with DHX36 signal decrease (Figure 3i). Of note, early binding of FANCJ at the G4 region was G4-structure independent and likely caused by binding to the non-G4 strand as shown previously^10^. Finally, RNaseA treatment retained DHX36 at the G4 region and prevented FANCJ from accumulating (Figure 3i). In conclusion, G-loop formation induces G4 unwinding through the helicase switch from DHX36 to FANCJ on the G4^G3N^, thereby allowing immediate incision after SLX4-XPF-ERCC1 recruitment.

## A general mechanism for G4 suppression

Finally, we addressed the generality of this G-loop-dependent G4 suppression mechanism. In our templates the displaced non-G4 strand contains a poly(T) motif that allows hybridization with the poly(A) tails from RNA transcripts present in extract (Figure S1b). However, the physiological counterpart likely contains a non-poly(T) heterogenous sequence. Therefore, we placed a 21-nt unique sequence across from the G4^G3N^ structure (pG4^BOT^-I, Figure S1b), and assessed G-loop formation in extract supplemented with an RNA transcript that contains a sequence complementary to this unique sequence (Figure 4a, Figure S6a,b). Importantly, while the transcript was slowly degraded over time in extract, we added it at a concentration that ensured a near 1:1 ratio of transcript to plasmid template at the time G-loop formation was expected to peak (Figure S6c,d). Addition of the complementary transcript to pG4^BOT^-I in extract promoted marked G-loop formation with a similar kinetics observed for pG4^BOT^ (Figure 4b). G-loop formation depended on homology between RNA and the non-G4 strand, because a non-homologous RNA transcript (transcript II) hardly promoted G-loop formation (Figure 4b). Conversely, a plasmid template that contained a sequence complementary to transcript II across from the G4 (pG4^BOT^-II) did induce G-loop formation (Figure S6e). Furthermore, both transcripts promoted G4 conversion in a homology-dependent fashion (Figure 4c, Figure S6f). Strikingly, while increasing the RNA transcript to plasmid ratio enhanced G-loop formation, it did not further enhance G4 conversion (Figure 4b,c, Figure S6e,f). Rather, it reduced G4 conversion, indicating that transcript abundance is an important regulatory variable in this process. In summary, DNA:RNA hybrids formed by complementary RNA transcripts across from G4s act in a general mechanism to suppress G4 structures.

**Figure 4.**
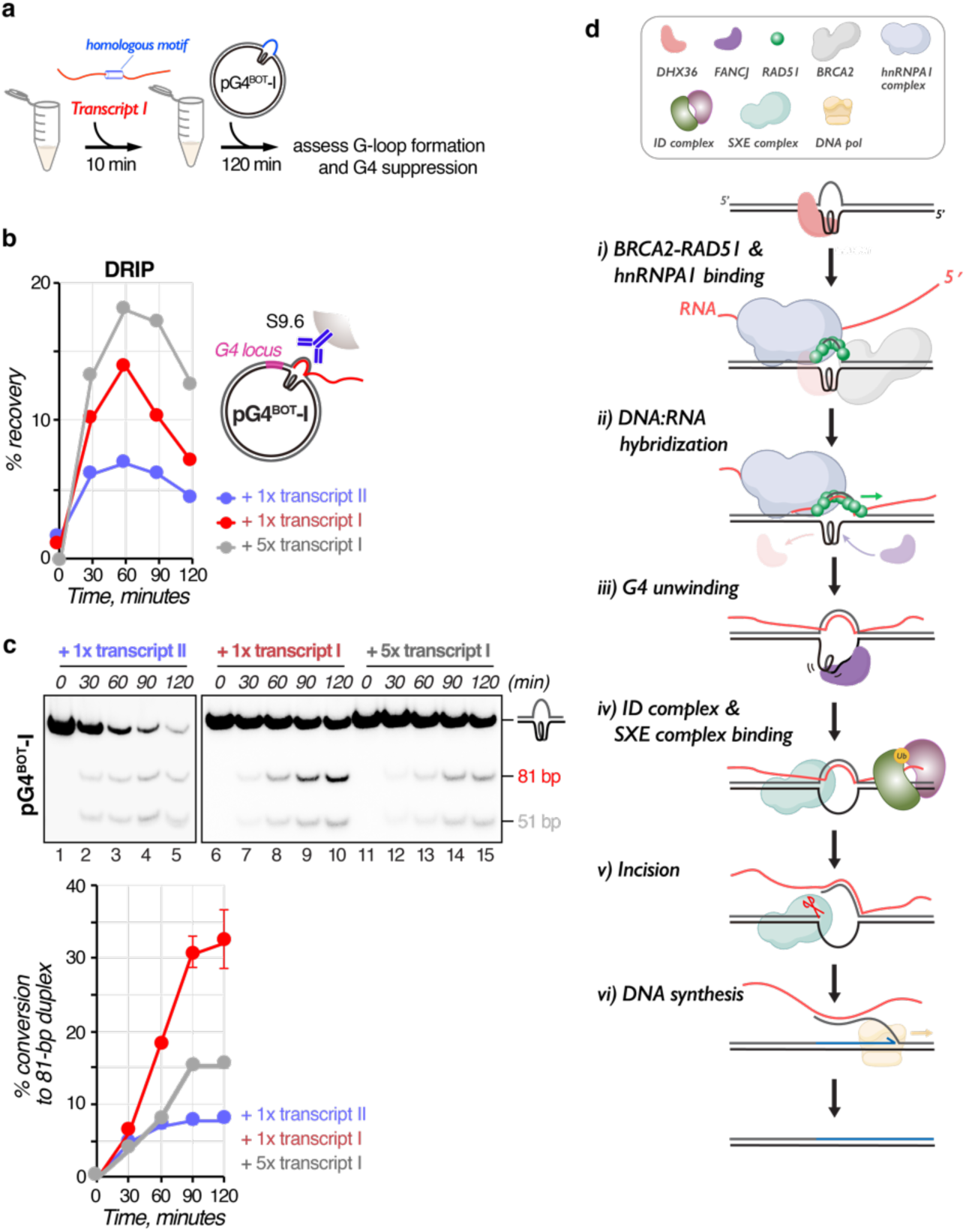
G-loop-dependent G-quadruplex suppression is a general mechanism. **a**, Experimental setup for the addition of synthesized RNA transcripts to G4-containing substrates with a sequence homologous to the RNA opposite the G4 (blue). DRIP and G4 conversion assays are performed on products generated upon incubation in NPE using these conditions. **b**, pG4^BOT^-I was incubated in NPE supplemented with transcript I or transcript II at 10 nM (1x) or 50 nM (5x) concentration, and products were analysed by DRIP-qPCR using primers for the G4 locus (schematic, right). Relative values compared to input signals were plotted. **c**, pG4^BOT^-I was incubated in NPE supplemented with transcript I or transcript II at indicated ratio, and products were isolated at various time points, digested with NotI and ClaI, end-labeled with ^32^P-α-dCTP, separated by native PAGE, and visualized by autoradiography (top). The 81-bp fragment was quantified and the conversion percentage was calculated by comparing their intensity to the total intensity of all fragments at time point 0. Conversion (%) was plotted against time with standard deviations (bottom, n=3). **d**, Model for R-loop-dependent G4 suppression.

## Discussion

Our data establish a versatile G4 suppression mechanism that acts through invasion of homologous RNA transcripts independently of ongoing transcription. DNA:RNA hybrid formation strictly depends on the presence of a G4 structure that provides a landing pad for HR factors and the hnRNPA1 complex, resulting in the formation of a G-loop (Figure 4d, i,ii). This promotes FANCJ loading followed by G4 unfolding and concomitant recruitment of the ID complex (Figure 4d, iii, iv). ID complex ubiquitylation then leads to the recruitment of the SLX4-XPF-ERCC1 nuclease complex that specifically incises the non-G4 strand to allow DNA synthesis past the G4 motif (Figure 4d, iv-vi). A tight connection of DNA:RNA hybrid formation and incisions allows rapid resolution of both G4 and R-loop structures to minimize potential collisions with DNA or RNA polymerases.

In contrast to a model that involves G4 unwinding and reannealing of the two strands, this G4 suppression mechanism is a highly regulated process involving nucleolytic incision and re-synthesis of the G4-opposing strand. Notably, incision of the non-G4 strand is accomplished by a mechanism reminiscent of ICL unhooking in the Fanconi anemia pathway^26^. This could explain severe telomere attrition observed in hematopoietic cells derived from Fanconi anemia patients^28^. Because we only observed specific incisions on the 5’ side of the non-G4 strand, DNA synthesis past the G4 most likely promotes strand displacement leading to a 5’ flap that is eventually removed by a flap endonuclease, as also observed during long-patch base excision repair. While this incision and re-synthesis mechanism may seem wasteful, we hypothesize that it could have important biological roles. First, the displaced ssDNA opposite a G4 structure is vulnerable to nucleolytic attack and cytosine deamination by APOBEC proteins, which could lead to DNA damage and mutations (Figure S7a). Replacement of this strand could therefore be important to maintain genome integrity. In line with this, the mutation frequency in the displaced non-G4 strand is sustained low despite enrichment of cytosine deaminase target motifs^29^. Second, it could be important for maintaining accessible chromatin at actively transcribed genes by erasing DNA methylations and removing nucleosomes (Figure S7b). Consistent with this, both G4 and R-loop structures are enriched in nucleosome-depleted and hypomethylated active promoters^14, 30–33^. Third, because DNA secondary structures including G4s form a barrier for the DNA replication machinery they are more likely to be left unreplicated upon entering mitosis. Indeed, G4 stabilization enhances mitotic DNA synthesis (MiDAS), indicating that G4-dependent incisions as described here may induce DSBs that trigger MiDAS^34^ in unreplicated G4-containing regions (Figure S7c). This is supported by our finding that G4 suppression is highly active in a mitotic *Xenopus* egg extract and recent reports describing that incision factors such as FANCD2 and SLX4 are required for MiDAS^35^.

Although R-loops are mostly described co-transcriptionally, they also stably form at sites lacking ongoing transcription, at which they overlap with G4 structures^4^. This supports the notion that R-loops and G4 structures function in a common mechanism that is not necessarily coupled to transcription. One important example of R-loop formation *in trans* is the invasion of repeat-containing RNAs (TERRAs) in mammalian telomeres which induces alternative lengthening of telomeres or ALT^36^. It is thought that RAD51 binds to TERRAs to promote strand exchange with donor dsDNA^15^. In contrast, in the mechanism we define, RAD51 is recruited to the displaced non-G4 DNA strand by BRCA2 and promotes homology-dependent hybridization with RNA transcripts shuttled by the hnRNPA1 complex (Figure 4d,ii). This enables efficient R-loop formation compared to strand exchange that RAD51 barely mediates *in vitro*^15^ and protects the displaced non-G4 strand from degradation, while ensuring RNA invasion exactly across from the G4 structure.

In addition to transcription-independent G4 suppression, we envision that a part of this mechanism may also be used to process co-transcriptional R-loops at sites where they overlap with G4s to prevent replisome collision that causes DNA damage (Figure S7d). Consistently, FANCD2, the FA core complex, and XPF-ERCC1 suppress co-transcriptional R-loops in mammalian cells^37, 38^. Because both G4 structures and co-transcriptional R-loops are elevated upon oncogenic stimuli^39, 40^, this mechanism may play a role in accelerated inflammation responses in cancer cells by generating DNA:RNA hybrid fragments that stimulate the cGAS-STING pathway after nuclear export^38^.

Overall, we uncover the first comprehensive mechanism that suppresses global G4 structures and G-loops and define previously uncharacterized functions for DNA repair enzymes linked to a broad range of human diseases. This provides insights into common pathologies such as cellular senescence and genome instability underlying the diseases. Inhibition of this mechanism by targeting the proteins or manipulating transcript levels can be exploited to develop therapeutic strategies for age-related diseases and cancer.

## Supporting information

Supplementary Figures, Tables, References

## Methods

### Preparation of plasmids

Plasmids containing a G4 motif were prepared as previously described^10^. Briefly, oligo DNA duplexes were generated by heating a pair of oligonucleotides (Table S1,2) at 80°C for 5 minutes and slowly cooling down to room temperature in ∼2 hours in annealing buffer (10 mM Tris-HCl pH 7.5 and 25 mM KCl). The resulting duplexes were ligated into the BbsI sites of the pSVR*lacO* vector^41^. After ligation, the closed circular plasmid was purified using a cesium chloride gradient ultracentrifugation, followed by butanol extraction, concentration and buffer exchange to TE (10 mM Tris-HCl pH7.5 and 1 mM EDTA) with an Amicon Ultra-4 centrifuge filter unit^42^ (30 kDa cut-off, Merck). Concentration of all plasmids was measured by NanoDrop One (Thermo Scientific) and the plasmids were stored at −80°C.

### Preparation of RNA transcripts

Transcript I and transcript II were synthesized from the pETDuet-1-KS1 and pETDuet-1-KS2 plasmids, respectively, at 37°C for 4 hours using the MEGAscript T7 transcription kit (Thermo Fisher Scientific) (see Table S3 for RNA sequences). To prevent read-through of T7 RNA polymerase, the plasmids were linearized with XhoI. For RNA transcript labelling, the reaction was supplemented with ^32^P-α-UTP. Concentration of the resulting RNA transcripts was measured by NanoDrop One, and purity of the transcripts was analysed by 4% urea-polyacrylamide gel prepared in 1x TBE buffer (89 mM tris-borate and 2 mM EDTA). The RNA transcripts were snap-frozen and stored at −80°C.

### Preparation of Xenopus egg extracts

*Xenopus laevis* female frogs (aged >2 years purchased from Nasco) were used as a source of eggs. High-speed supernatant (HSS), mitotic extract, demembranated sperm chromatin and nucleoplasmic extract (NPE) were prepared as previously described^43, 44^. All animal procedures and experiments were performed in accordance with national animal welfare laws and were reviewed by the Animal Ethics Committee of the Royal Netherlands Academy of Arts and Sciences (KNAW). All animal experiments were conducted under a project license granted by the Central Committee Animal Experimentation (CCD) of the Dutch government and approved by the Hubrecht Institute Animal Welfare Body (IvD), with project license number AVD80100201711044. Sample sizes were chosen based on previous experience, randomization and blinding are not relevant to this study.

### G-quadruplex suppression

G4 structures were induced on plasmids (75 ng/μL final concentration) prior to the G4 suppression reaction by incubating the plasmids at 80°C for 5 minutes and cooling down to room temperature in annealing buffer. Where indicated, the G4 ligand PhenDC_3_ (5 μM final concentration) was added to the plasmid DNA when it reached 50°C, the mixture was incubated at that temperature for 30 minutes, after which it was further cooled down to room temperature. To initiate G4 suppression upon DNA replication, the plasmids (9 ng/μL final concentration) were first incubated in HSS for 20 minutes at room temperature^45^. Two volumes of NPE (diluted to 40% with ELBS buffer containing 10 mM HEPES-KOH pH 7.7, 50 mM KCl, 2.5 mM MgCl_2_, and 250 mM sucrose) supplemented with 10 mM dithiothreitol, 15.5 mM phosphocreatine, 1.5 mM ATP, and creatine phosphokinase (3.8 ng/μl) were then added to start DNA replication. To monitor DNA synthesis, NPE was supplemented with ^32^P-α-dCTP. To inhibit DNA replication, recombinant *xl*Geminin was added to HSS at a 400 nM final concentration.

To study G4 suppression in NPE alone, the plasmids (9.3 ng/μL final concentration) were directly added to NPE diluted with ELBS buffer, supplemented with 2.5 mM dithiothreitol, 3.9 mM phosphocreatine, 0.39 mM ATP, and 1.0 ng/μL creatine phosphokinase, and incubated at room temperature. Where indicated, undiluted HSS or mitotic extract supplemented with 2.5 mM dithiothreitol, 3.9 mM phosphocreatine, 0.39 mM ATP, 1.0 ng/μL creatine phosphokinase and 1.3 ng/μL nocodazole was used for the reaction. Unless otherwise indicated, NPE diluted to 10% was used for the reaction. Where indicated, NPE was pre-incubated with RNaseA (1.25 mg/mL final concentration), RNaseH (1.0 unit/μL final concentration, New England Biolabs), α-amanitin (10 μM final concentration, Merck), transcript I, or transcript II (10 nM or 50 nM final concentration for each transcript) at room temperature for 30 minutes (for RNaseA and RNaseH) or 10 minutes (for α-amanitin and RNA transcripts) prior to reaction initiation by addition of plasmid DNA (t=0). For ChIP and DRIP experiments, an unrelated duplex control plasmid (pQuant, 0.44 ng/μL final concentration) was co-incubated to be used as an internal control for quantifications. At the indicated time, aliquots of the reaction (5 μL) were stopped with 45 μL stop solution II (50 mM Tris pH 7.5, 0.5% SDS, and 10 mM EDTA pH 8.0). Samples were then treated with RNaseA (0.15 mg/mL final concentration) for 30 minutes at 37°C, followed by Proteinase K (0.5 mg/mL final concentration) treatment at room temperature overnight. DNA was phenol/chloroform extracted, ethanol precipitated with glycogen (20 μg), and resuspended in 5 μL TE for further analysis.

### Incision assay

Extracted samples were digested with AflIII (0.004-0.006 unit/μL final concentration) at 37°C for one hour in 5 μL Buffer 3.1 (100 mM NaCl, 50 mM Tris-HCl, 10 mM MgCl_2_, 100 µg/mL bovine serum albumin (BSA), pH 7.9), end-labelled at 37°C for 30 minutes by addition of 1 μL end-labelling mix (0.125 mM dGTP, 0.125 mM dATP, 0.125 mM dTTP, 0.125 mM ^32^P-α-dCTP, Thermo Sequenase with thermostable Pyrophosphatase (Thermo Fisher), and 500 mM dithiothreitol), and mixed with 6 μL Gel Loading Buffer II. The samples were heated at 98°C for 5 minutes, snap-cooled on ice for 5 minutes, and immediately separated on a 7% urea-polyacrylamide sequencing gel prepared in 0.8x TTE buffer (71 mM Tris, 23 mM taurine, and 0.4 mM EDTA, pH 8.9). After gel drying, the products were visualized by autoradiography using an Amersham Typhoon (Cytiva). Sequencing ladders were generated using *primer T* and either pG4^TOP^ or pTEL^TOP^ with the Thermo Sequenase Cycle Sequencing kit (Thermo Fisher). The size of the incision products in pG4^TOP^ was determined using sequencing ladders and PCR-amplified “0” and “+1” fragments that were generated with pdsPolyT^BOT^, *primer T* labelled with ^32^P at the 5’-end and either *primer T-0-1* or *primer T-1-1* (see Table S2 for sequences). The size of the incision products in pTEL^TOP^ was determined by the same method with pdsPolyT^BOT^-II, *primer T* labelled with ^32^P at the 5’-end and either *primer T-0-2* or *primer T-1-2* (see Table S2 for sequences).

### G4 conversion assay

Extracted samples (1 μL) were digested with NotI-HF (0.89 unit/μL, New England Biolabs) and ClaI (0.44 unit/μL, New England Biolabs) in 9 μL CutSmart buffer (50 mM Potassium acetate, 20 mM Tris-acetate, 10 mM Magnesium acetate, 100 µg/ml BSA, pH 7.9), and end-labelled by the same method as described for the incision assay. The labeled samples were separated on a 12% polyacrylamide gel electrophoresis in 1× TBE buffer. After drying the gel with cellophane sheets (Thermo Fisher Scientific), products were quantified using an Amersham Typhoon and ImageQuant TL software (Cytiva). G4 conversion is plotted as the percentage of total intensity (measured at the time point 0) set to 100.

### DNA synthesis analysis

Extracted samples (3 μL) were digested with AflIII and BamHI (0.5 unit/μL and 1 unit/μL, respectively) at 37°C for 3 hours in 10 μL Buffer 3.1 and separated on a 1.4% agarose gel in 1× TBE buffer. After gel drying, the products were visualized by autoradiography and quantified using Amersham Typhoon and ImageQuant TL software. Nucleotide incorporation is plotted as the percentage of peak value with the highest value set to 100.

### RNA stability assay

Labelled transcript I and transcript II (10 nM final concentration for each) were incubated with 10% NPE at room temperature for 10 minutes before addition of homologous plasmid (pG4^BOT^-I or pG4^BOT^-II) to initiate G4 suppression. Aliquots of the reaction (5 μL) were stopped with 45 μL stop solution II supplemented with 1x RNAsecure (Thermo Fisher Scientific) and deproteinized with Proteinase K (0.2 mg/mL final concentration) at 37°C for one hour. RNAs were then purified with GeneJET RNA Cleanup and Concentration Micro Kit (Thermo Fisher Scientific). 8 μL eluate was mixed with 8 μL Gel Loading Buffer II, incubated at 98°C for 5 minutes, snap-cooled on ice for 5 minutes, and immediately separated on a 4% urea-polyacrylamide gel prepared in 1x TBE buffer. RNAs were visualized by SYBR Gold staining (Thermo Fisher Scientific) and autoradiography, and quantified using Amersham Typhoon and ImageQuant TL software (Cytiva). Intensities of labelled RNA transcripts were plotted as the percentage of value before incubation with NPE (the −10 minutes time point).

### Antibodies and immunodepletion

Antibodies against *xl*BRCA2^46^*, xl*DHX36^10^*, xl*FANCD2^47^*, xl*FANCJ^48^, *xl*MUS81^26^, *hs*RAD51^49^, *xl*SLX1^50^, *xl*SLX4^50^, *xl*XPA^51^, *xl*XPF^26^ and *hs*RNA Polymerase II subunit A (Bethyl) were previously described. Antibodies used for ChIP experiments were purified with rProtein A Sepharose (PAS) beads (Cytiva). The affinity-purified *xl*FANCA, and *xl*hnRNPA1 antibodies were raised against C-terminal residues (1406-1422: CSFKAPDDYDDLFFEPVF), and N-terminal residues (1-15: MHKSEAP(N/K)EPEQLRC), respectively by Vivitide.

For immunodepletion of proteins except for *xl*XPF and *xl*FANCA, one volume of Dynabeads Protein A beads (Thermo Fisher Scientific) was saturated with half volumes of antiserum or purified antibody (45 μg for 100 μL beads) by incubation for 30 minutes at room temperature. For mock depletion, pre-immunized rabbit serum or PAS-purified control rabbit IgG (Thermo Fisher Scientific) was used. To deplete NPE, one to three volume of each antibody-bound beads was incubated with one-half volumes of extract at room temperature for 30 minutes for two rounds (Table S4).

*xl*XPF and *xl*FANCA were depleted using the TRIM-away system^52^. Affinity-purified antibody against XPF (0.13 mg/mL final concentration) or FANCA (0.07 mg/mL final concentration) was mixed with NPE at 4°C for 15 minutes, recombinant human TRIM21 (0.2 mg/mL final concentration) was subsequently added, and the mixture was incubated at room temperature for 75 minutes.

### Protein purification

Recombinant N-terminal His_6_-tagged *xl*DHX36^10^, C-terminal FLAG-tagged *xl*FANCJ^48^, and *xl*SLX4 containing a N-terminal FLAG-tag and a C-terminal Strep-tag^50^, N-terminal FLAG-tagged *xl*XPF-C-terminal His_6_-tagged *hs*ERCC1^26^, and N-terminal His_6_-tagged *hs*TRIM21^52^ were prepared as previously described.

For purification of *xl*RAD51, the cDNA encoding wild-type *xl*RAD51 or *xl*RAD51^T128P^ (gBlocks Gene Fragments, Integrated DNA Technologies) was ligated into the NdeI-BamHI sites of the pET-15b vector (Merck). The wild-type protein was overexpressed as a N-terminal His_6_-tagged protein in the *E.coli* JM109(DE3) cells (Intact Genomics) cultured in 4.4 L LB medium and purified by the previously described method^53^, in which the His_6_-tag was removed with the thrombin protease (Cytiva). *xl*RAD51^T128P^ was purified as the wild-type protein but without spermidine precipitation. The proteins were snap-frozen and stored at −80°C. For rescue experiments, pG4^BOT^ was incubated with purified xlRAD51 or *xl*RAD51^T128P^ (9 μM final concentration) in binding buffer containing 20 mM HEPES pH 7.4, 1 mM dithiothreitol, 2.5 mM MgCl_2_, and 1 mM ATP at room temperature for 20 minutes before NPE addition.

For purification of the *xl*hnRNPA1 complex, 700 μL NPE was mixed with Dynabeads Protein A beads bound with *xl*hnRNPA1 antibody at room temperature for 30 minutes. After removing supernatant, proteins were eluted in buffer A (20 mM Tris-HCl pH9.0, 10% glycerol, 50 mM NaCl, and 2 mM dithiothreitol) containing the *xl*hnRNPA1 N-terminal peptide (0.38 mg/mL final concentration) by incubating at 4°C for 30 minutes. The eluate was applied to a Mono Q 5/50 GL (pre-equilibrated in buffer A) (Cytiva), and washed with 10 mL buffer A. *xl*hnRNPA1 complex was eluted with a 10 mL linear gradient of 50 to 500 mM NaCl in buffer A. Fractions that reversed G4 suppression deficiency by hnRNPA1 depletion (fractions between 6 mL to 8 mL in elution volume) were collected and then loaded to Superose 6 Increase 10/300 GL (Cytiva) that was pre-equilibrated with ELBS buffer containing 2 mM dithiothreitol. Fractions between 15.7 mL to 19.2 mL in elution volume were concentrated with Amicon Ultra-4 Centrifugal Filter Unit (10 kDa cutoff, Merck) to ∼2 mg/mL (measured by Bradford method), snap-frozen, and stored at −80°C. For rescue experiments, this purified hnRNPA1 complex (∼1 mg/mL final concentration) is incubated with NPE at room temperature for 10 minutes before plasmid addition. The concentration of all recombinant proteins was determined by SDS-PAGE with Coomassie Brilliant Blue staining, using BSA as a standard protein.

### DNA:RNA Immunoprecipitation (DRIP)

Extracted DNA (4.5 μL) was incubated with the S9.6 antibody (2.0 μg, Sigma-Aldrich) in 100 μL DRIP buffer (10 mM sodium phosphate pH 7.0, 9 mM Tris-HCl pH7.5, 0.9 mM EDTA, 140 mM NaCl, and 0.05% Triton X-100) at 4°C overnight. Where indicated, extracted DNA was first digested with AflIII (0.5 unit/μL final concentration, New England Biolabs) and BamHI (1 unit/μL final concentration, New England Biolabs) in 10 μL buffer 3.1 or treated with with RNaseH (0.25 unit/μL final concentration) in 10 µL RNaseH buffer (50 mM Tris-HCl, 75 mM KCl, 3 mM MgCl_2_, 10 mM dithiothreitol, pH 8.3) at 37°C for 2 hours prior to the antibody addition. To capture the DNA-antibody complex, 15 μL Dynabeads Protein A beads (pre-equilibrated with DRIP buffer) were added, incubated on a rotating wheel at 4°C for 2 hours, and then washed with 375 µL DRIP buffer twice at room temperature for 15 minutes. Captured DNA was released by deproteinization in 153 µL DRIP elution buffer (50 mM Tris pH8.0, 10 mM EDTA, 0.5% SDS, and 0.11 mg/mL Proteinase K) at 55°C for 45 minutes. DNA was phenol/chloroform-extracted, dissolved in 30 μL 10 mM Tris-HCl pH7.5, and analyzed by quantitative PCR in 10 µL reaction buffer (6 mM Tris pH 8.3, 25 mM KCl, 2.5 mM MgCl_2_, 0.3 mM dNTPs, 0.1% Tween-20, 0.1 mg/ml BSA, 1:66,500 SYBR Green I (Sigma-Aldrich), and Hot Start Taq DNA polymerase), using 0.25 µM of following primer pairs (Table S1): *G4 for* and *G4 rev* (for *G4* locus, 37-136 bp upstream from G4s), *lacO for* and *lacO rev* (for *lacO* locus, 295-388 bp downstream from G4s), and *pQuant for* and *pQuant rev* (for assessment of background binding of the proteins on pQuant). The values from *pQuant* primers were subtracted from the values for *G4* and *lacO* primers.

### Chromatin immunoprecipitation (ChIP)

ChIP was performed similar to described previously^54^. At the indicated times, replication samples (3 μL) were crosslinked with 47 μL ELBS buffer containing 1% formaldehyde at room temperature for 10 minutes. After quenching the formaldehyde by addition of 5 μL 1.25 M glycine, the samples were passed through a Micro Bio-Spin 6 Chromatography column (Bio-Rad), added to 445 μL Sonication buffer (20 mM Tris-HCl pH7.5, 150 mM NaCl, 2 mM EDTA, 0.5% IGEPAL CA-630, 2 mM phenylmethylsulfonyl fluoride, 5 μg/mL aprotinin, and 5 μg/mL leupeptin), and sonicated. Sonicated samples (30 μL) were incubated with the indicated antibodies (5 μg) at 4°C overnight, and the antibody complex was captured by mixing with 10 μL PAS beads at room temperature for 2 hours. The protein-bound DNA fragments were washed extensively with 475 μL Sonication buffer once, with 475 μL Sonication buffer containing 500 mM NaCl and 100 mM KCl twice, with 475 μL ChIP wash buffer (10 mM Tris-HCl pH7.5, 250 mM LiCl, 1 mM EDTA, 0.5% IGEPAL CA-630, and 0.5% SDS) twice, and then with 475 μL TE once. The DNA fragments were eluted in ChIP elution buffer (50 mM Tris pH7.5, 10 mM EDTA, and 1% SDS) by incubating at 65°C for 20 minutes. After treatment with RNaseA (0.1 mg/mL final concentration) at 37°C for 30 minutes, the crosslinked proteins were digested and reversed by consecutive incubation at 42°C for 6 hours and then at 70°C for 9 hours with 275 mM NaCl and 0.12 mg/mL Proteinase K. The released DNA was phenol/chloroform-extracted, followed by quantitative PCR by the same method as described for DRIP. The values from *pQuant* primers were subtracted from the values for *G4* primers. ChIP data are plotted as the percentage of peak value with the highest value set to 100.

## Acknowledgements

We thank R. Kanaar, F. Mattiroli, S. Noordermeer, M. Takata, A. N. Zelensky for feedback on the manuscript, the Hubrecht animal caretakers for animal support, and the other members of the Knipscheer laboratory for discussions. We thank K. Cimprich for the XPA antibody, R. Kanaar for the RAD51 antibody, S. Noordermeer for initial help with the DRIP-qPCR protocol, and R. de Boer for the mitotic extract. K.S. was supported by the Kanae Foundation for the Promotion of Medical Science and Osamu Hayaishi Memorial Scholarship for Study Abroad from the Japanese Biochemical Society. This work was supported the European research council (ERC) through an ERC consolidator grant (ERCCOG 101003210-XlinkRepair) to PK and the Gravitation program CancerGenomiCs.nl from the Netherlands Organisation for Scientific Research (NWO). PK is also supported by the Oncode Institute, which is partly financed by the Dutch Cancer Society (KWF).

## Author Contributions

K.S. designed the study, performed all the experiments, analysed data and wrote the manuscript. A.G.M.H developed and established critical experimental methods. P.K. designed and supervised the study and wrote the manuscript.

## Competing interests

Authors declare no competing interests.

## Notes

### Competing Interest Statement

The authors have declared no competing interest.

