## Supplementary Figures, Tables, References for "RNA transcripts suppress G-quadruplex structures through G-loop formation"

Supplementary Figure S1-7

Supplementary Table S1-4

Supplementary References

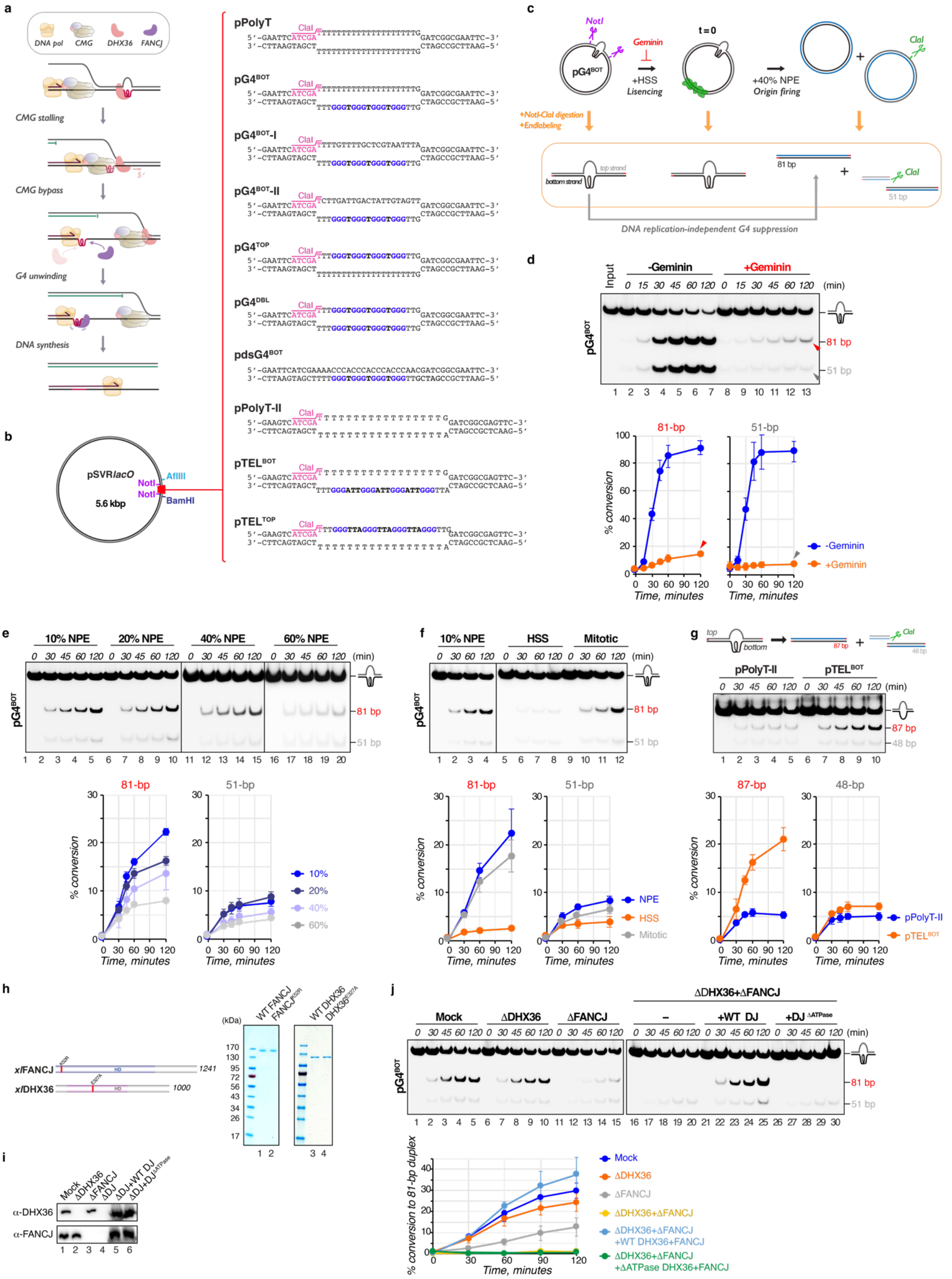
a
DNA pol
CMG
DHX36
FANCJ
CMG stalling
CMG bypass
G4 unwinding
DNA synthesis
pPolyT
pG4BOT
pG4BOT-I
pG4BOT-II
pG4TOP
pG4DBL
pdsG4BOT
pPolyT-II
pTELBOT
pTELTOP
ClaI
b
pSVRlacO
5.6 kbp
NotI
AflIII
BamHI
c
NotI
pG4BOT
Geminin
+HSS
Licensing
t = 0
+40% NPE
Origin firing
+NotI-ClaI digestion
+Endlabeling
top strand
bottom strand
81 bp
51 bp
DNA replication-independent G4 suppression
d
Input
-Geminin
+Geminin
(min)
81 bp
51 bp
81-bp
51-bp
% conversion
Time, minutes
e
10% NPE
20% NPE
40% NPE
60% NPE
f
HSS
Mitotic
NPE
g
top
bottom
87 bp
48 bp
pPolyT-II
pTELBOT
87-bp
48-bp
h
xlFANCJ
xlDHX36
HD
1241
1000
(kDa)
WT FANCJ
FANCJK52R
WT DHX36
DHX36E327A
i
Mock
ΔDHX36
ΔFANCJ
ΔDJ
ΔDJ+WT DJ
ΔDJ+DJΔATPase
α-DHX36
α-FANCJ
j
ΔDHX36+ΔFANCJ
+WT DJ
+DJ ΔATPase
% conversion to 81-bp duplex
ΔDHX36+ΔFANCJ
+WT DHX36+FANCJ
+ΔATPase DHX36+FANCJ

**Figure S1 | *Xenopus* egg extracts support replication-independent G-quadruplex suppression**

**a,** Model for DNA replication-coupled G4 suppression. DHX36 binds to a G4 with high affinity[55] before the replication machinery arrives at the structure. Stalling of the CMG helicase at the G4 stimulates translocation of DHX36, generating ssDNA past the G4. The ssDNA promotes CMG bypass of the intact G4 structure and allows the leading strand to approach to the G4. This second stalling event leads to the recruitment of FANCJ to the G4 and triggers G4 unfolding, allowing the leading strand to extend past the G4 motif. When DHX36 is absent, FANCJ promotes the CMG bypass by translocating on the non-G4 strand past the G4 structure. When FANCJ is absent, DHX36 re-accumulates at the G4s and unwinds the structures. This redundancy confers robustness to this pathway.

**b,** Schematic representation and sequence information of plasmids used in this study. Inserted (non)complementary duplex sequences containing a G4 or a control motif are indicated (right). A unique ClaI restriction site (magenta) and G-tracts (blue) are indicated. Restriction sites used in this study are indicated on the plasmid (left). Because Watson and Crick base pairing in dsDNA is more favorable than G4 Hoogsteen base pairing, G4 structures are preferentially formed in ssDNA[56]. Therefore, a non-complementary sequence across from the G4 motif is necessary to stably form G4 structures in these plasmids.

**c,** Scheme of products generated before and after pG4$^{BOT}$ replication followed by digestion with NotI (purple) and ClaI (green) and subsequent end-labeling. DNA replication in egg extract involves incubation in High-speed supernatant of egg lysate (HSS) to allow the assembly of pre-replication complexes (Pre-RC, green ovals), followed by addition of a nucleoplasmic extract (NPE) to trigger origin firing and a single round of DNA replication. ClaI specifically digests the replicated top strand, generating a 51-bp product, while the replicated bottom strand that contains no ClaI motif is detected as an 81-bp product. Addition of Geminin to HSS prevents Pre-RC assembly, thereby blocking DNA replication initiation.

**d,** pG4$^{BOT}$ was replicated in egg extract in the presence or absence of Geminin. At various times (t = 0 is immediately after NPE addition. Input represents the products prior to HSS incubation.) DNA was isolated, digested with NotI and ClaI, end-labeled with $^{32}$P-$\alpha$-dCTP, separated by native PAGE, and visualized by autoradiography (top). The 81-bp and 51-bp fragments were quantified and the conversion percentage was calculated by comparing their intensity to the total intensity of all fragments at time point 0.  Conversion (%) was plotted against time with standard deviations (bottom, n=3). Where indicated, HSS was supplemented with Geminin to block DNA replication initiation. In the presence of Geminin, the 81-bp and 51-bp fragments both still slowly accumulated with a preference for the 81-bp, suggesting the existence of DNA replication-independent G4 suppression mechanism in *Xenopus* egg extracts (**c**, gray arrow).

**e,** pG4$^{BOT}$ was incubated in NPE (diluted to 60, 40, 20, or 10%) and products were analysed as in **d** (n=3). Since NPE contains a high concentration of Geminin, no DNA replication initiates when the plasmid is incubated directly in NPE[57]. While incubation with 10% NPE yielded ~20% of the 81-bp product with a strong preference over the 51-bp product, both the yield and preference were compromised at increasing concentrations of NPE, suggesting that the replication-independent G4 suppression is active under specific concentrations of its regulatory factor(s).

**f,** pG4$^{BOT}$ was incubated in NPE, HSS, or mitotic extract (Mitotic) and products were analysed as in **d** (n=3). In HSS, hardly any 81-bp or 51-pb product was generated and the preference for the 81-bp product was lost. In 10% NPE or mitotic extract, the 81-bp was preferably generated and increased till the last time point while less of the 51-bp product was formed and this hardly accumulated at later times. These results indicates that the 81-bp product is formed, at least in part, by a different mechanism.

**g,** pPolyT-II and pTEL$^{BOT}$ were incubated in NPE, and products were analysed as in **d** (top). The 87-bp (corresponding to the 81-bp product in pG4$^{BOT}$) and 48-bp (corresponding to the 51-bp product in pG4$^{BOT}$) products were quantified as in **d** (n=3). As G4$^{TEL}$ forms different G4 structures compared to G4$^{G3N}$ [58], G4 suppression in *Xenopus* egg extracts acts on diverse G4 structures. Minor dsDNA fragments also accumulate in pPolyT with different kinetics, indicating that both the G4 and the non-G4 strand are also converted to duplex DNA by a G4-independent background mechanism.

**h**, Recombinant purified *Xenopus laevis* FANCJ and DHX36 proteins used in this study were analysed by SDS-PAGE with Coomassie Brilliant Blue staining alongside molecular-weight

markers (right). Schematic representation of both proteins with their domain organization is shown (left). Mutated residues are indicated with red bars. HD, helicase domain.
**i**, Mock-, DHX36-, FANCJ-, and DHX36-FANCJ-depleted NPEs supplemented with buffer, wild-type DHX36 and FANCJ, or ATPase-dead DHX36 and FANCJ mutants were analysed by Western blot with DHX36 and FANCJ antibodies.
**j**, pG4$^{BOT}$ was incubated in the NPEs as described in **I**, and products were analysed as in **d**.

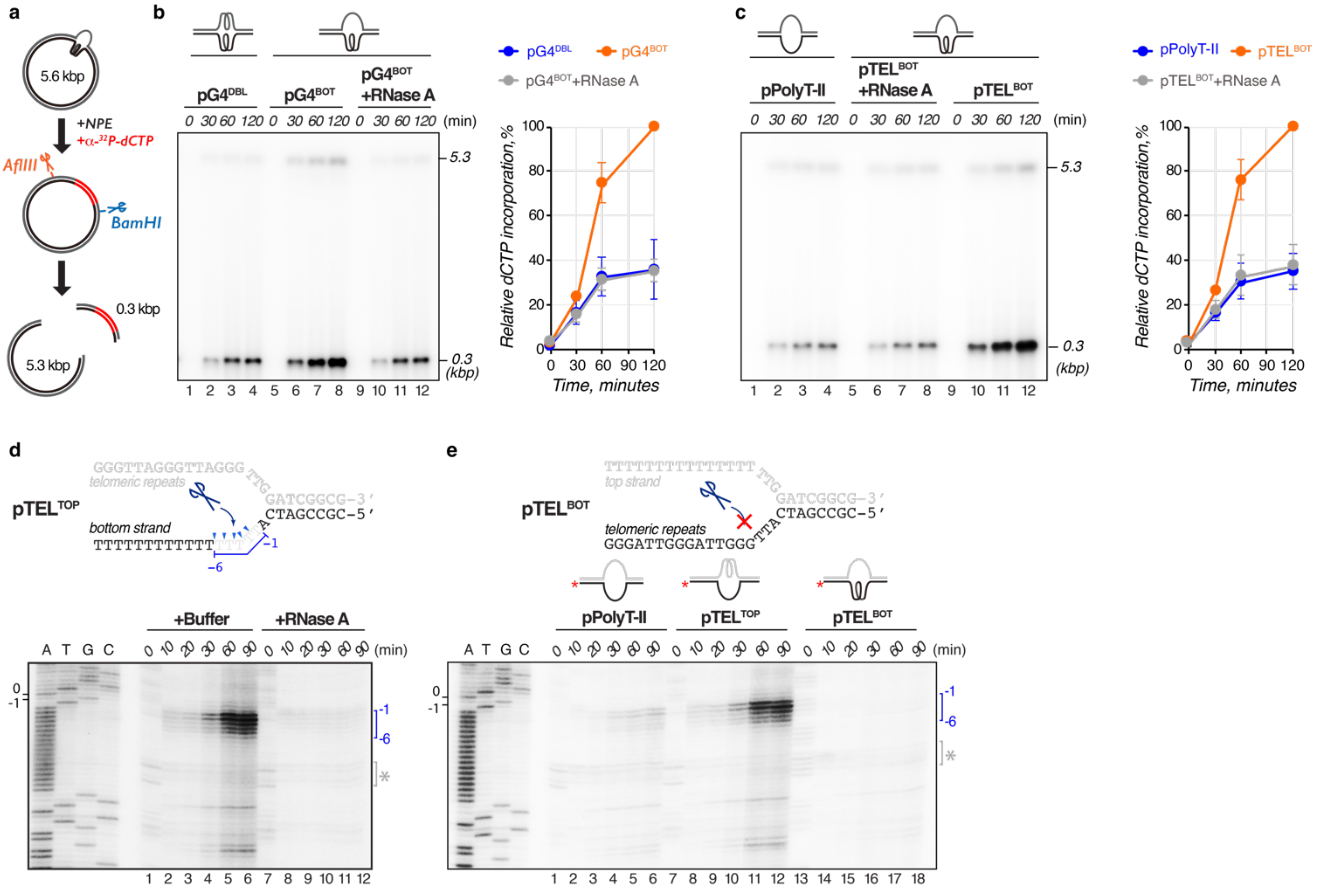


**Figure S2 | G-quadruplex suppression involves nucleolytic incision and DNA synthesis**

**a**,**b**, pG4$^{BOT}$ and pG4$^{DBL}$ were incubated in NPE supplemented with $^{32}$P-α-dCTP. At various times, DNA was isolated, digested with AflIII and BamHI, separated on an agarose gel, and visualized by autoradiography (Schematic, **a**). Double digestion with AflIII and BamHI excises a ~270-bp fragment containing the non-complementary region from the plasmid backbone (~5.3 kbp). $^{32}$P-α-dCTP incorporation in the ~270-bp fragment was quantified and the relative intensities compared to the highest signal among the conditions were plotted with standard deviations (**b**, right, n=3). Where indicated, NPE was pre-treated with RNaseA. While $^{32}$P-α-dCTP was preferentially incorporated in the ~270-bp region of pG4$^{BOT}$ after 30 minutes, the incorporation was compromised when NPE is pre-treated with RNaseA or when the G4 is present on both strands (pG4$^{DBL}$). These results indicate that efficient DNA synthesis requires both an unstructured DNA strand across from the G4 structure and endogenous RNA transcripts.

**c**, pPolyT-II and pTEL$^{BOT}$ were incubated in NPE supplemented with $^{32}$P-α-dCTP, and products were analysed as in **b** (n=3). Similar to the result in **b**, $^{32}$P-α-dCTP incorporation in the ~270-bp fragment of pTEL$^{BOT}$ was compromised when NPE is pre-treated with RNaseA or when the G4 was omitted (pPolyT-II). This indicates that efficient DNA synthesis requires both a G4 structure and endogenous RNA transcripts.

**d**, pTEL$^{TOP}$ was incubated in NPE pre-treated with RNaseA or buffer. At various times, DNA was isolated, digested with AflIII, end-labeled with $^{32}$P-α-dCTP, separated by denaturing PAGE alongside a sequencing ladder, and visualized by autoradiography. Sequence surrounding the incision sites of pTEL$^{TOP}$ is depicted and incision positions are indicated (top, blue arrow heads). Incision products are numbered -6 to -1, where the -1 product corresponds to a product in which incision takes place between the first and the second nucleotides upstream of the ss/dsDNA junction. A bracket with an asterisk indicates products generated by star activity of AflIII.

**e**, pPolyT-II, pTEL$^{TOP}$, and pTEL$^{BOT}$ were incubated in NPE, and products were analysed by denaturing PAGE as in **d**. While incubation of pTEL$^{TOP}$ yielded the -1 to -6 products, these products were hardly observed when G4$^{TEL}$ was situated on the bottom strand (pTEL$^{BOT}$) or omitted (pPolyT-II). These results indicate that the non-G4 strand is site-specifically incised and this depends on the presence of the G4 structure. Therefore, the low-level conversion observed on both strands in pPolyT-II (Figure S1g) is most likely mediated by non-specific nucleolytic attacks on the ssDNA. The incised ends seem to be processed by extensive resection in pPolyT-II, resulting in loss of NotI motif(s), since the intensity of the fragment containing the non-complementary region in pPolyT-II decreased over time (Figure S1g). Similar decrease was observed with pPolyT (Figure 1a). Interestingly, this was hardly observed in G4-containing plasmids, suggesting that a G4 structure stimulates a protective mechanism(s).

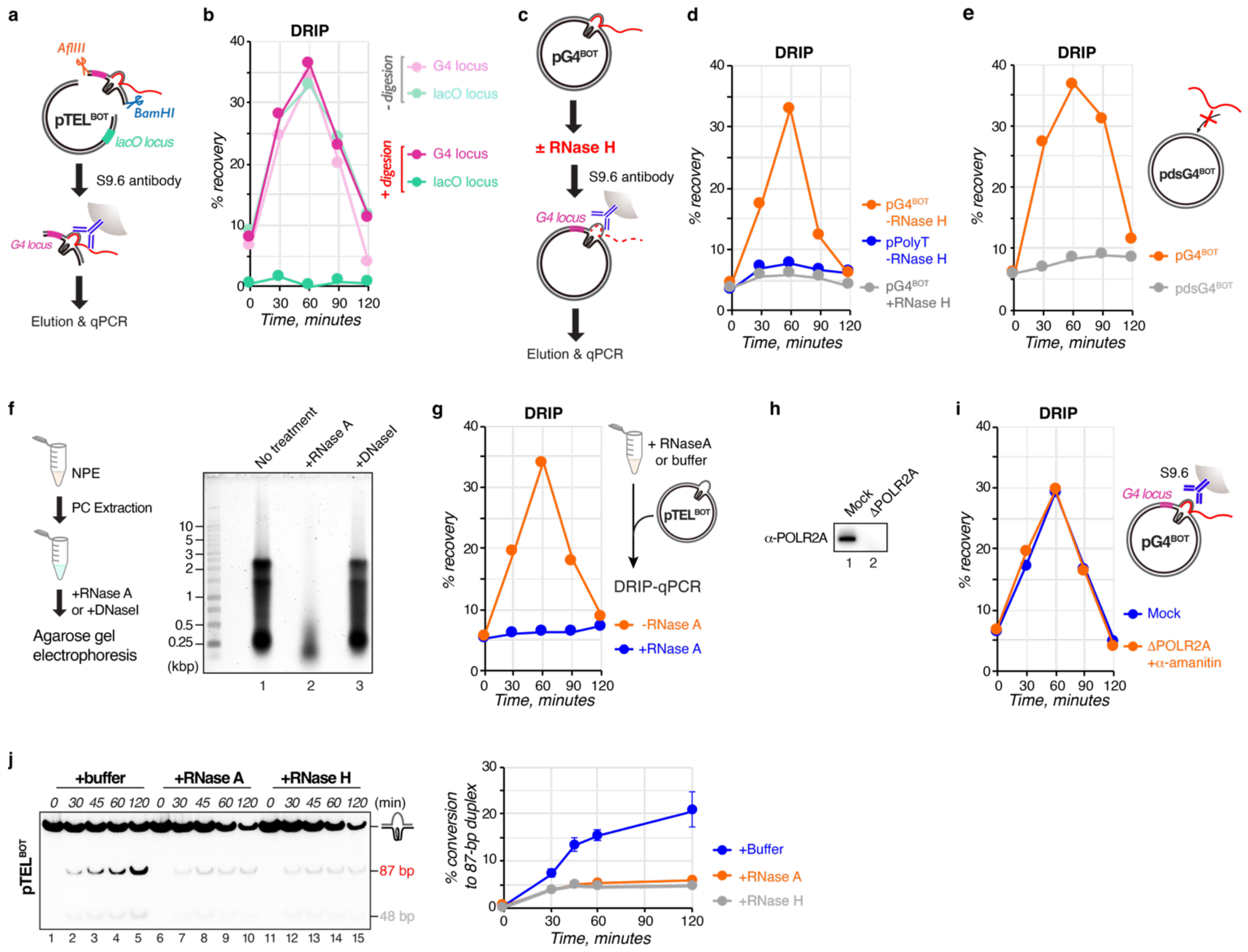

**Figure S3 | RNA transcripts hybridize with a displaced strand across from a G-quadruplex independently of active transcription.**

**a**,**b**, pTEL$^{BOT}$ was incubated in NPE, and products were isolated, when indicated digested with AflIII and BamHI, and immunoprecipitated with the S9.6 antibody (blue, left schematic, **a**). The co-precipitated DNA was amplified by quantitative PCR (qPCR) with primers specific to the G4 locus (magenta) or the *lacO* locus (green, left schematic, **a**). Relative values compared to input signals for each locus were plotted (**b**). AflIII and BamHI digestion, which excised a ~270-bp fragment containing the non-complementary region, prior to immunoprecipitation abolished signals at the *lacO* locus while retaining G4 locus signal. This indicates that DNA:RNA hybrids are specifically formed at the G4 containing fragment.

**c**, Schematic of DRIP-qPCR assay used in **d**.

**d,** pPolyT and pG4$^{BOT}$ were incubated in NPE, and products were isolated, treated with RNaseH or buffer, and analysed by DRIP-qPCR as in **b** with primers for the G4 locus. Relative values compared to input signals were plotted. Pre-treatment with RNaseH abolished signals in pG4$^{BOT}$, confirming that this method specifically detects DNA:RNA hybrids.

**e**, pG4$^{BOT}$ and pdsG4$^{BOT}$ were incubated in NPE, and products were analysed by DRIP-qPCR as in **b**. When the G4 motif was situated in dsDNA, DNA:RNA hybrids hardly accumulated, indicating that DNA:RNA hybrid formation depends on formation of the G4 structure.

**f**, Nucleic acids were extracted from NPE with phenol-chloroform (PC), digested with RNaseA or DNaseI, or remained untreated. Products were analysed by agarose gel electrophoresis with SYBR-Gold staining alongside a DNA ladder. Roughly 2 μg nucleic acids were extracted from 1 μL NPE. The SYBR-Gold staining intensity was highly sensitive to RNaseA but not DNaseI, indicating that NPE contains high concentration of RNA transcripts. Consistent with this, the extracted nucleic acids exhibited a ratio of 260 nm/280 nm absorbance of 2, which indicates that they consist of RNA. Since *Xenopus* eggs are inactive in transcription[59], the RNA transcripts are most likely derived from the oocytes.

**g**, pTEL$^{BOT}$ was incubated in NPE that was pre-treated with RNaseA or buffer, and products were analysed by DRIP-qPCR as in **d**.

**h,** Mock-depleted and RNA polymerase II subunit A (POLR2A)-depleted NPEs were analysed by Western blot with a POLR2A antibody.

**i,** pG4$^{BOT}$ was incubated in the NPEs as described in **h**, and products were analysed by DRIP-qPCR as in **d**. Where indicated, NPE was supplemented with $\alpha$-amanitin, an inhibitor of eukaryotic RNA polymerases.

**j,** pTEL$^{BOT}$ was incubated in NPE that was pre-treated with RNaseA, RNaseH, or buffer. At various times, DNA was isolated, digested with NotI and ClaI, end-labeled with $^{32}$P-$\alpha$-dCTP, separated by native PAGE, and visualized by autoradiography (top). The 87-bp fragment was quantified and the conversion percentage was calculated by comparing its intensity to the total intensity of all fragments at time point 0. Conversion (%) was plotted against time with standard deviations (bottom, n=3). In the RNaseA- and RNaseH-treated extracts, the intensity of the fragment containing the non-complementary region markedly declined over time, which is reminiscent of processive degradation of a non-complementary region observed in both pPolyT and pPolyT-II. Similar decrease was observed for pG4$^{BOT}$ (Figure 1g). These data indicate that the DNA:RNA hybrids protect the G4-forming regions from degradation.

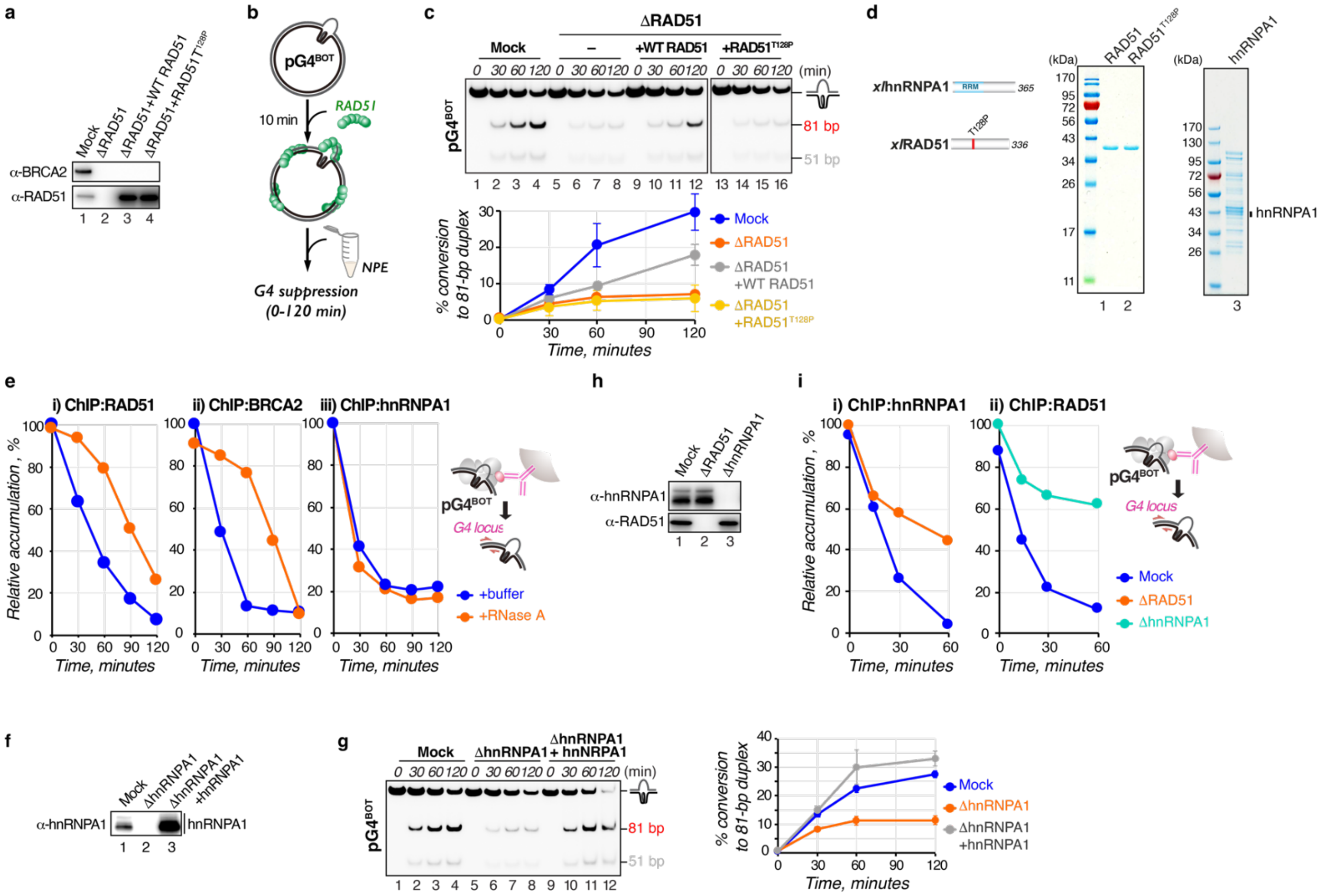


**Figure S4 | RAD51 and the hnRNPA1 complex collaborate to form a G-loop to suppress G-quadruplexes**

**a**, Mock- and RAD51-depleted NPEs, where indicated supplemented with wild-type (WT) RAD51, or RAD51$^{T128P}$, were analysed by Western blot with RAD51 and BRCA2 antibodies. RAD51 depletion efficiently codepletes BRCA2.

**b**, **c**, pG4$^{BOT}$ was incubated with buffer, WT RAD51, or RAD51$^{T128P}$ to allow pre-binding of the RAD51 proteins (schematic, **b**) and subsequently incubated in the NPEs as described in **a**. At various times, DNA was isolated, digested with NotI and ClaI, end-labeled with $^{32}$P-$\alpha$-dCTP, separated by native PAGE, and visualized by autoradiography (top). The 81-bp fragment was quantified and the conversion percentage was calculated by comparing its intensity to the total intensity of all fragments at time point 0. Conversion (%) was plotted against time with standard deviations (bottom, n=3).

**d**, Purified *Xenopus laevis* (*xl*) RAD51 and the hnRNPA1 complex isolated and purified from NPE were analysed by SDS-PAGE with Coomassie Brilliant Blue staining alongside molecular-weight markers (right). Schematic representation of both proteins with their domain organization is shown (left). The mutated residue in RAD51 is indicated with a red bar. RRM, RNA recognition motif.

**e**, pG4$^{BOT}$ was incubated in NPE that was pre-treated with RnaseA or buffer, and products were analysed by ChIP-qPCR with (i) RAD51, (ii) BRCA2, and (iii) hnRNPA1 antibodies using primers for the G4 locus. The relative values compared to the highest signal among the conditions were plotted.

**f**, Mock- and hnRNPA1-depleted NPEs where indicated supplemented the hnRNPA1 complex were analysed by Western blot with a hnRNPA1 antibody.

**g**, pG4$^{BOT}$ was incubated in the NPEs as described in **f**, and products were analysed as in **c** (n=3).

**h**, Mock-, RAD51-, and hnRNPA1-depleted NPEs were analysed by Western blot with RAD51 and hnRNPA1 antibodies.

**i,** pG4$^{BOT}$ was incubated in the NPEs as described in **h**, and products were analysed by ChIP-qPCR as in **e** with hnRNPA1(i) and RAD51 (ii) antibodies.

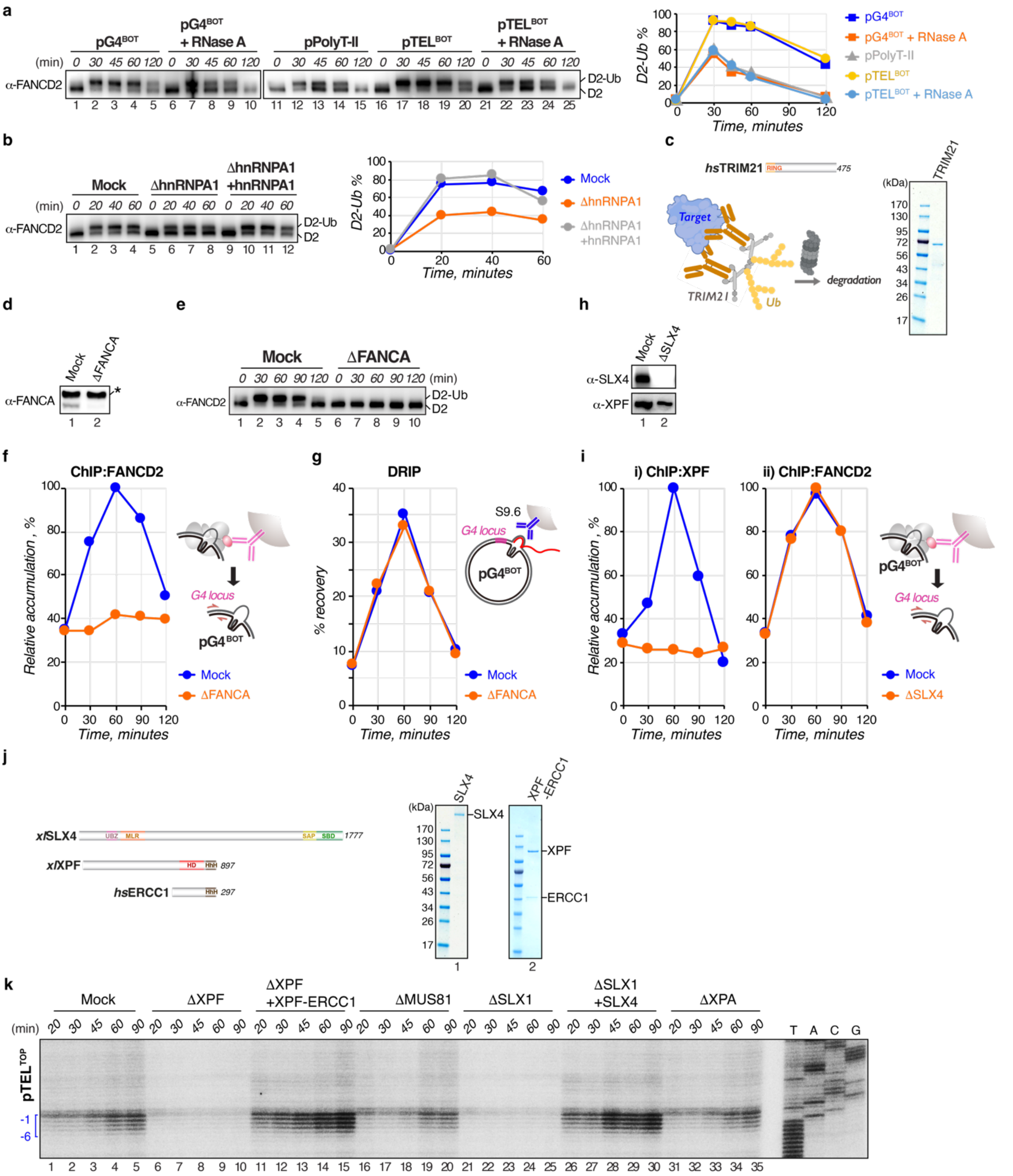


**Figure S5 | G-loops recruit the SLX4-XPF-ERCC1 nuclease complex by inducing FANCD2 monoubiquitination**

**a,** pG4$^{BOT}$, pPolyT-II, and pTEL$^{BOT}$ were incubated in NPE and the extract was analysed at various times by Western blotting with a FANCD2 antibody. Where indicated, NPE was pre-treated with RNaseA. The intensity of monoubiquitinated and unmodified FANCD2 (D2-Ub and D2, respectively) were quantified and the D2-Ub/D2 ratio were plotted (right). While FANCD2 was robustly monoubiquitinated in both pG4$^{BOT}$ and pTEL$^{BOT}$, the modification was compromised when the G4 structures are omitted or after RNaseA treatment, indicating that FANCD2 ubiquitination is promoted by the G4 structure and DNA:RNA hybrid. Residual FANCD2 modification is most likely

due to non-specific binding of the ID complex to the plasmid backbone that stimulates FANCD2 monoubiquitination[60].
**b**, pG4$^{BOT}$ was incubated in the NPEs as described in Figure 2e and analysed by Western blotting with a FANCD2 antibody. D2-Ub/D2 ratio were plotted as described in **a** (right).
**c**, Purified *Homo sapiens* (*hs*) TRIM21 protein used in this study was analysed by SDS-PAGE with Coomassie Brilliant Blue staining alongside molecular-weight markers (right). Schematic representation of the protein with a domain organization is shown together with a schematic of Trim-Away-mediated protein degradation[52] (left). Extract is first incubated with an antibody against a target protein to allow antibody-antigen binding. Subsequent addition of hsTRIM21 induces acute and rapid proteasome-mediated degradation of the target protein, antibody and TRIM21 complex. RING, RING Finger domain.
**d**, FANCA was depleted from NPE using the Trim-away system described in **c**. Mock-depleted and FANCA-depleted NPEs were analysed by Western blot with the FANCA antibody. An asterisk represents a non-specific band.
**e**, pG4$^{BOT}$ was incubated in the NPEs as described in **d**, and the extracts were analysed at various times by Western blotting with a FANCD2 antibody.
**f**, pG4$^{BOT}$ was incubated in the NPEs as described in **d**, and products were analysed by ChIP-qPCR with a FANCD2 antibody using primers for the G4 locus. The relative values compared to the highest signal among the conditions were plotted.
**g**, pG4$^{BOT}$ was incubated in the NPEs as described in **d**, and products were analysed by DRIP-qPCR with primers for the G4 locus. Relative values compared to input signals were plotted.
**h**, Mock- and SLX4-depleted NPEs were analysed by Western blot with SLX4 and XPF antibodies.
**i**, pG4$^{BOT}$ was incubated in the NPEs as described in **h**, and products were analysed by ChIP-qPCR as in **f** with XPF (i) and FANCD2 (ii) antibodies. SLX4 depletion prevented XPF accumulation at the G4 locus, while retaining FANCD2 accumulation. This indicates that XPF is recruited as a complex with SLX4 downstream of FANCD2 accumulation.
**j**, Purified *Xenopus laevis* (*xl*) SLX4 and the *xl*XPF-*hs*ERCC1 complex used in this study were analysed by SDS-PAGE with Coomassie Brilliant Blue staining alongside molecular-weight markers (right). Schematic representation of the proteins with a domain organization is shown (left). UBZ, ubiquitin binding domain; MLR, MUS312/MEI-9 interaction like region; BTB, broad complex-tram-track-bric-a-brac domain; SAP, SAF-A/B-Acinus and PAIS domain; SBD, SLX1-binding domain; HD, helicase domain; nuclease domain; HhH, helix-hairpin-helix.
**k**, pTEL$^{TOP}$ was incubated in the NPEs as described in Figure 3d, and products were digested with AflIII, end-labelled, separated by denaturing PAGE alongside a sequencing ladder, and visualized by autoradiography. Incised fragments (-1 to -6) are indicated with a bracket. Site-specific incision in pTEL$^{TOP}$ was fully dependent on SLX4-XPF-ERCC1 as observed for pG4$^{TOP}$ (Figure 3e). Consistent with these observations, XPF-, or ERCC1-depeted cells are sensitive to a G4 ligand while MUS81- or XPA-depleted cells are not[61]. Among the incision products, only the -1 product is consistent with *in vitro* activity of the SLX4-XPF-ERCC1 complex[62]. Since the shorter products (up to -8 for pG4$^{TOP}$ and -6 for pTEL$^{TOP}$) accumulate over time, we speculate that site-specific incisions on a displaced strand consist of two steps; incision at -1 by the SLX4-XPF-ERCC1 and subsequent 5' to 3' resection. Further resection might be blocked by DNA:RNA hybrid[63] or proteins bound to the ssDNA.

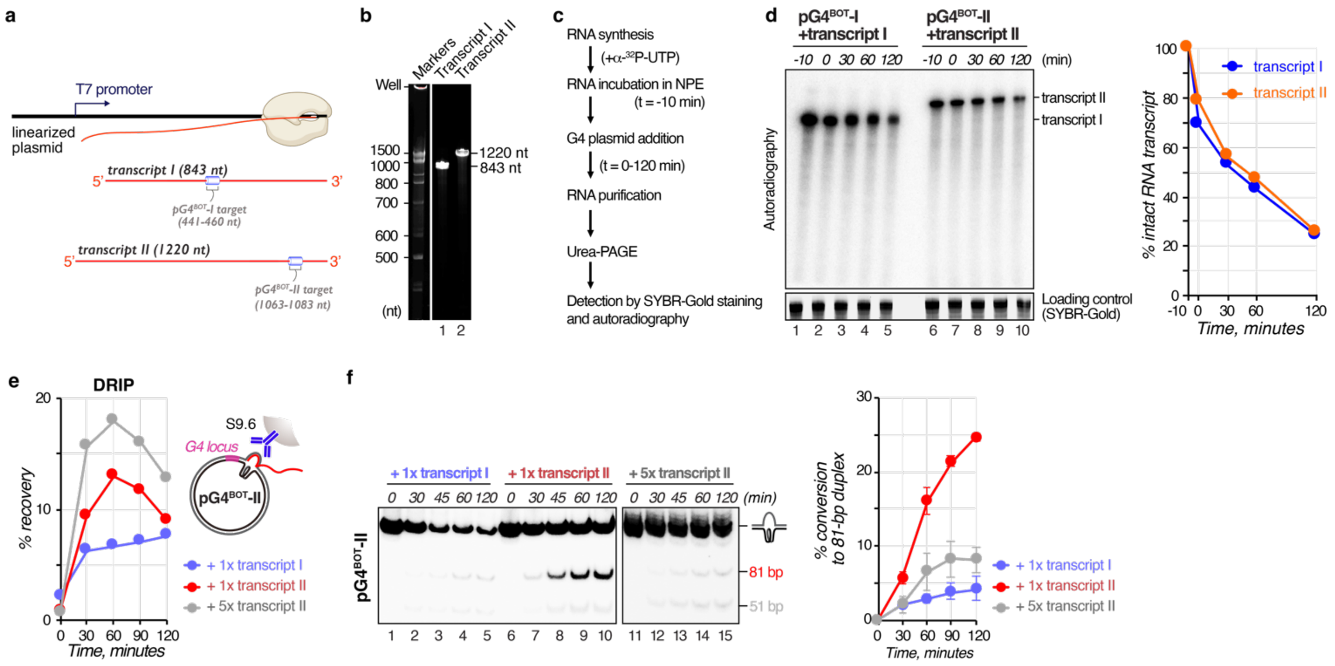


**Figure S6 | G-quadruplex suppression depends on RNA transcripts complementary to a displaced strand across from the structure**

**a,** Schematic representation of RNA transcripts used in this study. The RNAs were synthesized by T7 RNA polymerase from a T7 promoter in linearized plasmids encoding either transcript I (843 nt) or transcript II (1,220 nt) (Table S3 for sequences). Complementary regions to the displaced non-G4 sequence in pG4$^{BOT}$-I or pG4$^{BOT}$-II are indicated with brackets.

**b**, Synthesized transcript I and transcript II were analysed by denaturing PAGE with ethidium bromide staining alongside a DNA ladder.

**c,** Schematic diagram of the RNA stability assay. RNAs are synthesized in the presence of $^{32}$P-$\alpha$-UTP and incubated in NPE for 10 minutes to allow interaction with RNA binding factors. To initiate G4 suppression, a G4-containing plasmid with a non-G4 strand complementary to the RNAs (pG4$^{BOT}$-I or pG4$^{BOT}$-II) was added. At various time points, RNA was isolated, separated by denaturing PAGE, and visualized by SYBR-Gold staining and autoradiography.

**d**, Transcript I and transcript II labeled with $^{32}$P-$\alpha$-UTP were incubated in NPE and analyzed by the RNA stability assay (**c**). Intact transcript I and transcript II were quantified and the relative intensities compared to the signal at time point -10 (before incubation in NPE) were plotted (right). t = 0 is immediately after plasmid addition. Although the intensities of both transcripts declined, approximately 50% of the transcripts remained intact at 30 minutes when G-loop formation initiates. Endogenous RNA transcripts detected by SYBR-Gold staining are shown as a loading control.

**e**, pG4$^{BOT}$-II was incubated in NPE supplemented with transcript I or transcript II at 10 nM (1x) or 50 nM (5x) concentration, and products were analysed by DRIP-qPCR using primers for the G4 locus (schematic, right). Relative values compared to input signals were plotted. Transcript II, but not transcript I, promoted G-loop formation, indicating that G-loop formation depends on homology between RNA and the non-G4 strand.

**f**, pG4$^{BOT}$-II was incubated in NPE supplemented with transcript I or transcript II at 10 nM (1x) or 50 nM (5x) concentration, and products were isolated at various time points, digested with NotI and ClaI, end-labeled with $^{32}$P-$\alpha$-dCTP, separated by native PAGE, and visualized by autoradiography (top). The 81-bp fragment was quantified and the conversion percentage was calculated by comparing its intensity to the total intensity of all fragments at time point 0. Conversion (%) was plotted against time with standard deviations (bottom, n=3).

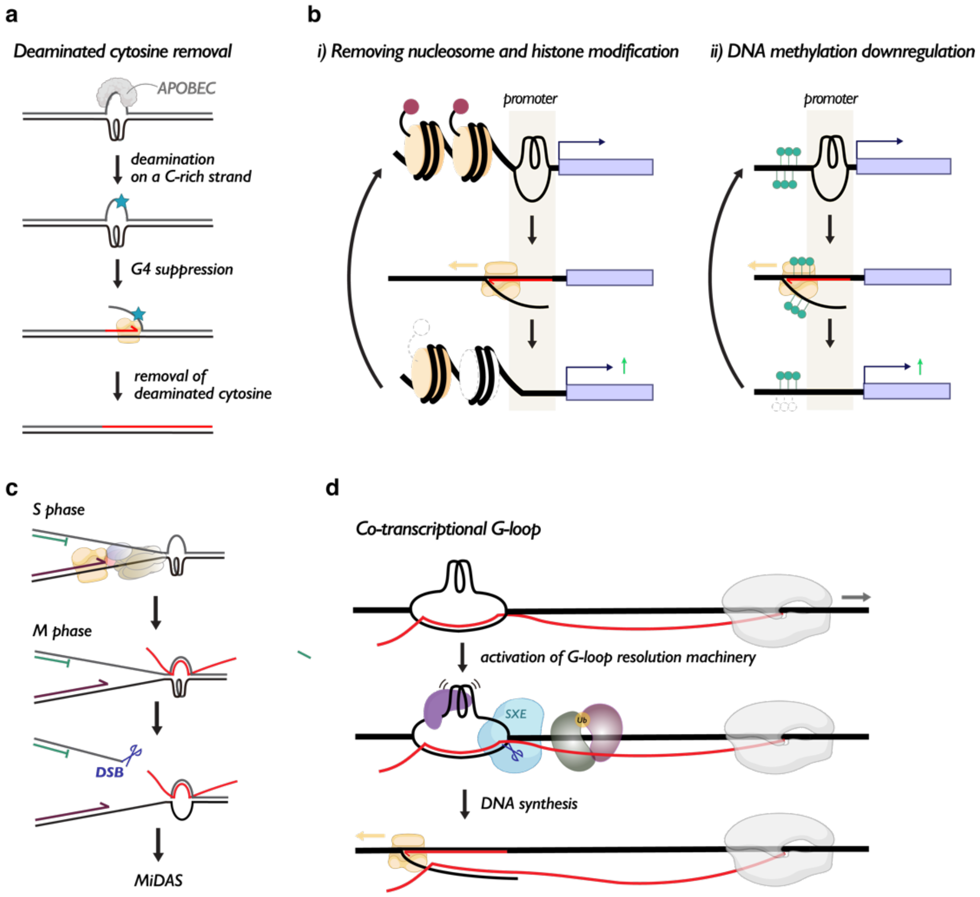


**Figure S7 | Possible biological roles of incision-dependent G4 suppression**

**a**, Model for deaminated cytosine removal. Displaced non-G4 sequences are vulnerable of apolipoprotein B mRNA editing cytosine deaminases (APOBECs)[64], which ultimately induces T:G mismatches. DNA synthesis past the G4 motif removes the deaminated cytosine (blue), preventing the mutational consequence.

**b**, Model for open chromatin establishment. DNA synthesis past the G4 motif can eliminate nucleosomes (i), epigenetic histone marks (i), and DNA methylation (ii) adjacent to the G4 structure. This is advantageous to establish and sustain transcriptionally active promoters. This could also be linked to synthesis-dependent histone variant deposition in promoters, establishing histone variant boundaries that define early replication origins[65]. G4 structures are indeed found enriched in active replication origins[66].

**c,** Model for MiDAS initiation through G4 suppression. When a replication fork stalls at a G4 structure, the unreplicated structure can remain in M phase. In mitosis, the G-loop mediated G4 suppression mechanism can induce a one-ended double-strand break that can stimulate break induced recombination that repairs the DSBs, known as mitotic DNA synthesis (MiDAS)[67].

**d,** Model for co-transcriptional G-loop resolution. A G-loop could be formed behind transcription in high GC skew regions where C-rich DNA forms a DNA:RNA hybrid with G-rich RNA[32]. Once formed, the G4 structure is resolved by DHX36 and FANCJ, while the DNA:RNA hybrid stimulates FANCD2 ubiquitination and subsequent recruitment of the SLX4-XPF-ERCC1 complex. Incision and DNA synthesis release the DNA:RNA hybrid from the template strand.

| *Oligo name* | *Sequence* |
|---|---|
| ***A*** | 5′-CCCTACAAGAATTCATCGATTTTTTTTTTTTTTTTTTTTGGATCGGCGAATTCCCCG-3′ |
| ***B*** | 5′-GCACCGGGGAATTCGCCGATCGTTTTTTTTTTTTTTTTTTTTCGATGAATTCTTGT-3′ |
| ***C*** | 5′-CCCTACAAGAATTCATCGATTTGGGTGGGTGGGTGGGTTGGATCGGCGAATTCCCCG-3′ |
| ***D*** | 5′-GCACCGGGGAATTCGCCGATCGTTGGGTGGGTGGGTGGGTTTCGATGAATTCTTGT-3′ |
| ***E*** | 5′-CCCTACAAGAATTCATCGATTTTGTTTTGCTCGTAATTTAGATCGGCGAATTCCCCG-3′ |
| ***F*** | 5′-CCCTACAAGAATTCATCGATCTTGATTGACTATTGTAGTTGATCGGCGAATTCCCCG-3′ |
| ***G*** | 5′-CCCTACAAGAATTCATCGAAAACCCACCCACCCACCCAACGATCGGCGAATTCCCCG-3′ |
| ***H*** | 5′-CCCTACAAGAAGTCATCGATTTTTTTTTTTTTTTTGGATCGGCGAGTTCCCCG-3′ |
| ***I*** | 5′-GCACCGGGGAACTCGCCGATCATTTTTTTTTTTTTTTTCGATGACTTCTTGT-3′ |
| ***J*** | 5′-CCCTACAAGAAGTCATCGATTTGGGTTAGGGTTAGGGTTAGGGTTGGATCGGCGAGTTCCCCG-3′ |
| ***K*** | 5′-GCACCGGGGAACTCGCCGATCATTGGGTTAGGGTTAGGGTTAGGGTTTCGATGACTTCTTGT-3′ |
| ***L*** | 5′-CCCTACAAGAATTCATCGAAAAAAAAAAAAAAAAAAAAACGATCGGCGAATTCCCCG-3′ |
| ***M*** | 5′-CCCTACAAGAAGTCATCGAAAAAAAAAAAAAAAAATGATCGGCGAGTTCCCCG-3′ |
| ***T*** | 5′-GATCCGCGCCCAATACGCAAACCGCCTC-3′ |
| ***G4 for*** | 5′-TTTTCCTCCTCTCCTGACTACTCCC-3′ |
| ***G4 rev*** | 5′-CATGCATTGGTTCTGCACTTCC-3′ |
| ***lacO for*** | 5′-AGCTAACTTGTTTATTGCGAATTCGG-3′ |
| ***lacO rev*** | 5′- TTTAGTGAGGGTTAATTGCGCGC-3′ |
| ***pQuant for*** | 5′-TACAAATGTACGGCCAGCAA-3′ |
| ***pQuant rev*** | 5′-GAGTATGAGGGAAGCGGTGA-3′ |
| ***T-0-1*** | 5′-GTTTTTTTTTTTTTTTTTTTTTCGATGAATTC-3′ |
| ***T-1-1*** | 5′-CGTTTTTTTTTTTTTTTTTTTTTCGATGAATTC-3′ |
| ***T-0-2*** | 5′-ATTTTTTTTTTTTTTTTTTCGATGACTTCTTG-3′ |
| ***T-1-2*** | 5′-CATTTTTTTTTTTTTTTTTTCGATGACTTCTTG-3′ |

**Table S1 |** Sequences of oligonucleotides used in this study.

| *plasmid* | *Oligonucleotide pair (top strand + bottom strand)* |
|---|---|
| pPolyT | *Oligo A + Oligo B* |
| pG4$^{TOP}$ | *Oligo C + Oligo B* |
| pG4$^{BOT}$ | *Oligo A + Oligo D* |
| pG4$^{BOT}$-1 | *Oligo E + Oligo D* |
| pG4$^{BOT}$-2 | *Oligo F + Oligo D* |
| pG4$^{DBL}$ | *Oligo C + Oligo D* |
| pdsG4$^{BOT}$ | *Oligo G + Oligo D* |
| pPolyT-2 | *Oligo H + Oligo I* |
| pTEL$^{TOP}$ | *Oligo J + Oligo I* |
| pTEL$^{BOT}$ | *Oligo H + Oligo K* |
| pdsPolyT$^{BOT}$ | *Oligo L + Oligo B* |
| pdsPolyT$^{BOT}$-2 | *Oligo M + Oligo I* |

**Table S2 |** Oligonucleotide pairs used for plasmid generation in this study.

| ***Transcript I (843 nt)*** |
|---|
| 5′–GGGGAAUUGUGAGCGGAUAACAAUUCCCCUCUAGAAAUAAUUUUGUUUAACUUUAAGAAGGAGAUAUACCAUGG<br>GCAGCAGCCAUCACCAUCAUCACCACAGCCAGGAUCCGUCGUUUAGUACAGCUUCUGGAAAAACAGUACAACUUUCGGA<br>CGAGUCGUUAAAACGCGCGCGUGCUUUUUUCUCGGAGAUCGACAAUUCCCAACUUUUACAACAACUGAAUAACGAGUCC<br>CCCUUUGAUGAACGCUCAAUCGGCGUGGAGAUGAAGAAGUCGAUUCAGACACCGUUGACCACAGAGAAGGCGGAAACAA<br>CGGAAUCCAUCAAGGUGAAAGUUAACAACGGGAGCUUUGGGUUUAAUACAACAUCUGGCAAACAGGUUUCGGUGAGUGA<br>GUCUGCGCUGAAGAAGGUGAAAGACAUUUUUCAGGAAUGUGAUGAUUCGGUAAAUUACGAGCAAAACAAACCCUUGGUU<br>CGUAUGACCAUGAGUUCAAAAAUGAAGGAAUCCGCUCCCGGGCUGAAGCGUCCUGCCCAAAUCCCCGUAUCCAGUUACC<br>AGUCCGACAAUGUCCAGAGCAAGGAAGGAAAUGUUAGUACAUUUCAGGACGGCUAUGCGAAUAAGAAAUCCUUAAAUAC<br>GUACUCAGAGGAGGCCCGCCGUCCUGCGAUCACAACUGUCCCCCGCCAACAUCUGACCCCCACGCAUAUCUCAAAGAUC<br>AGUACAUCAACGAGCGCCUUCCCCAACAAAUCAAAGCCUGAUUUGUACGUGACGACAUCACACAACACUCCGCAGAAUG<br>AUUUUGAGAUCGAGGCCGCGGAGUCUGCUCGUGCAUUCUUAGAAUAAUAAUAACUCGA–3′ |
| ***Transcript II (1220 nt)*** |
| 5′–GGGGAAUUGUGAGCGGAUAACAAUUCCCCUCUAGAAAUAAUUUUGUUUAACUUUAAGAAGGAGAUAUACCAUGG<br>GCAGCAGCCAUCACCAUCAUCACCACAGCCAGGAUCCAUGCACAAAUCUGAAGCCCCUAACGAGCCGGAGCAACUCCGC<br>AAACUCUUUAUCGGCGGUCUCUCCUUUGAGACGACGGACGAAUCACUCCGGGAGCACUUUGAGCAGUGGGGUACAUUAA<br>CAGAUUGCGUCGUAAUGCGGGACCCUAACUCCAAACGUAGCCGGGGGUUCGGUUUUGUAACUUACCUCAGCACUGAUGA<br>GGUUGACGCUGCCAUGACAGCACGUCCGCAUAAGGUGGAUGGUCGGGUGGUGGAGCCAAAACGUGCCGUCUCUCGGGAA<br>GACUCAUCGCGGCCAGGCGCUCAUCUCACCGUAAAAAAGAUUUUUGUCGGUGGUAUUAAAGAAGACACUGAGGAAGAUC<br>ACCUUCGCGAGUAUUUUGAACAAUACGGGAAAAUCGAGGUUAUUGAGAUCAUGACCGACCGCGGUAGCGGGAAGAAACG<br>UGGUUUUGCCUUUGUCACGUUUGAAGACCAUGACUCCGUCGACAAGAUUGUUAUUCAAAAGUACCAUACGGUAAACAAU<br>CAUAAUUGUGAGGUGCGCAAAGCGCUUUCCAAGCAAGAGAUGGCUAGCGUUUCUGGGAGCCAACGCGGCCGCGGGGGUU<br>CAGGUAACUAUGGGUCUCGUGGCGGGUUUGGGAACGACAACUUCGGUGGGCGCGGUGGGAACUUCGGGGGGAACCGUGG<br>GGGUGGGGGUGGGUUUGGCAACCGCGGGUAUGGGGGCGAUGGCUAUAAUGGGGAUGGCAACUAUGGUGGCAGCCCGCCA<br>UAUUCCGGCGGGAACCGUGGCUACGGGGCAGGUCAGGGUGGGGGCUACGGGGCGGGCCAAGGUGGUGGUUAUGGGGGUG<br>GUGGGCAAGGCGGUGGUUACGGGGGGAACGGGGGGUACGAUGGGUACAAUGGUGGCGGUUCAGGUUUCUCUGGUAGCGG<br>GGGCAACUUUGGCAGCUCAGGCGGGUAUAACGACUUUGGCAACUACAAUAGUCAAUCAAGUUCCAAUUUUGGUCCUAUG<br>AAAGGUGGCAAUUAUGGUGGUGGUCGUAACAGUGGUCCUUAUGGUGGUGGGUACGGUGGUGGCAGCGCCAGUAGCAGCU<br>CCGGGUAUGGCGGGGGCCGUCGCUUCUAAUAAUAACUCGA–3′ |

**Table S3 |** Sequences of RNA transcripts used in this study. Homologous sequences to pG4$^{BOT}$-I and pG4$^{BOT}$-II are indicated in blue.

| *Protein* | *Antibody* | *Ratio (Extract:beads)* | *Source of antibody* |
|---|---|---|---|
| POLR2A | *Affinity-purified* | **1:2.5** | *Bethyl, Cat#A300-653A* |
| RAD51 | *PAS-purified* | **1:1.5** | *Tan et al., 1999* |
| BRCA2 | *Anti-serum* | **1:1.5** | *Brandsma et al., 2019* |
| FANCD2 | *Affinity-purified* | **1:1.5** | *this study* |
| hnRNPA1 | *Affinity-purified* | **1:2** | *this study* |
| SLX4 | *Anti-serum* | **1:1.5** | *Hoogenboom et al., 2019* |
| SLX1 | *Anti-serum* | **1:1.5** | *Hoogenboom et al., 2019* |
| MUS81 | *Anti-serum* | **1:1.5** | *Klein Douwel et al., 2014* |
| XPA | *Anti-serum* | **1:1** | *Bomgarden et al., 2006* |
| DHX36 | *Anti-serum* | **1:0.67** | *Sato et al., 2021* |
| FANCJ | *Anti-serum* | **1:1.33** | *Castillo Bosch et al., 2014* |

**Table S4 |** Immunodepletion conditions used in this study.
